# Disentangling cause and consequence: Genetic dissection of the *DANGEROUS MIX2* risk locus, and activation of the DM2h NLR in autoimmunity

**DOI:** 10.1101/2020.11.01.363895

**Authors:** Jana Ordon, Patrick Martin, Jessica Lee Erickson, Filiz Ferik, Gerd Balcke, Ulla Bonas, Johannes Stuttmann

## Abstract

Nucleotide-binding domain–leucine-rich repeat-type immune receptors (NLRs) protect plants against pathogenic microbes through intracellular detection of effector proteins. However, this comes at a cost, as NLRs can also induce detrimental autoimmunity in genetic interactions with foreign alleles. This may occur when independently evolved genomes are combined in inter- or intraspecific crosses, or when foreign alleles are introduced by mutagenesis or transgenesis. Most autoimmunity-inducing NLRs are encoded within highly variable *NLR* gene clusters with no known immune functions, which were termed autoimmune risk loci. Whether risk NLRs differ from sensor NLRs operating in natural pathogen resistance and how risk NLRs are activated in autoimmunity is unknown. Here, we analyzed the *DANGEROUS MIX2* risk locus, a major autoimmunity hotspot in *Arabidopsis thaliana*. By gene editing and heterologous expression, we show that a single gene, *DM2h*, is necessary and sufficient for autoimmune induction in three independent cases of autoimmunity in accession Landsberg *erecta*. We focus on autoimmunity provoked by an EDS1-YFP^NLS^ fusion protein to functionally characterize DM2h and determine features of EDS1-YFP^NLS^ activating the immune receptor. Our data suggest that risk NLRs function reminiscent of sensor NLRs, while autoimmunity-inducing properties of EDS1-YFP^NLS^ are in this context unrelated to the protein’s functions as immune regulator. We propose that autoimmunity may, at least in some cases, be caused by spurious, stochastic interactions of foreign alleles with co-incidentally matching risk NLRs.

## Introduction

Plants growing in natural habitats, as well as crops cultivated in the field, are constantly exposed to pathogenic microbes, which can induce severe damage and yield losses. However, plants evolved two major types of immune receptors to protect from disease (Jones and Dangl, 2006; Dodds and Rathjen, 2010). Extracellularly, membrane-anchored pattern recognition receptors (PRRs) can detect microbe-associated molecular patterns (MAMPs), which induces a suite of responses collectively termed pattern-triggered immunity (PTI; e.g. reviewed in Saijo et al., 2018). Intracellularly, immune receptors belonging to the nucleotide-binding domain–leucine-rich repeat (NLR) class can detect microbial proteins secreted into host cells during the infection process, termed effector proteins, thus inducing effector-triggered immunity (ETI; *e.g*. reviewed in Cui et al., 2015). Although ETI and MTI are qualitatively similar, they differ in amplitude, and especially ETI is often accompanied by localized cell death at infection sites - the hypersensitive response (HR). So far, ETI and PTI have largely been considered as independent signaling networks, however, recent reports suggest that plant resistance depends on mutual amplification of PTI and ETI responses (Ngou et al., 2020; Yuan et al., 2020).

Plant NLRs are modular proteins consisting of three domains: A C-terminal leucine-rich repeat (LRR) domain, a central nucleotide-binding (NB-ARC) domain and either a coiled coil (CC) or a Toll/interleukin-1 receptor (TIR) domain at the N-terminus (Bentham et al., 2017; Song et al., 2020). These N-terminal domains initiate immune signaling upon receptor activation, and subdivide plant NLRs into TNLs (TIR domain), CNLs (CC domain) and RNLs, which hold an atypical CC domain with homology to the non-NLR resistance protein RESISTANCE TO POWDERY MILDEW8 (RPW8; Meyers et al., 2003; Collier et al., 2011; Jubic et al., 2019). Structurally similar immune receptors, albeit differing in their N-terminal domains, function in animals and fungi. Activation and signaling by animal NLRs involves oligomerization and interactor recruitment by oligomeric N-terminal assemblies (Bentham et al., 2017). Recent structural elucidation of a full length plant CNL (HOPZ-ACTIVATED RESISTANCE1 (ZAR1); Wang et al., 2019a; Wang et al., 2019b) and TNL (Recognition of XopQ1 (Roq1); Martin et al., 2020b) revealed assembly of pentameric and tetrameric resistosomes by the activated immune receptors, respectively. Resistosome formation is driven by indirect (ZAR1) or direct (Roq1) pathogen effector recognition, nucleotide exchange (ADP/ATP) within the nucleotide-binding domain and profound conformational changes. In the ZAR1 resistosome, the N-terminal CC domains assemble into a funnel-like structure, which was proposed to translocate into membranes. Thereby, the ZAR1 resistosome might induce the HR directly by interfering with membrane integrity or by functioning as a selective ion channel (Wang et al., 2019a). In the resistosome formed by activated Roq1, two dimers of TIR domain dimers exhibit a two-fold symmetry. This conformation leads to opening of the NADase active site of the TIR domain and breakdown of NAD^+^ (Horsefield et al., 2019; Wan et al., 2019; Duxbury et al., 2020; Martin et al., 2020b). In the latter case, it is assumed that one of the breakdown products of the TIR domain enzymatic activity, whose molecular identity remains to be determined, functions as a signaling molecule. In agreement with an indirect mode of action, resistance mediated by TNLs further requires proteins of the ENHANCED DISEASE SUSCEPTIBILITY1 (EDS1) family and RNL-type helper NLRs of the N requirement gene1 (NRG1) and/or the ACTIVATED DISEASE RESISTANCE1 (ADR1) class (Wagner et al., 2013; Castel et al., 2018; Wu et al., 2018; Gantner et al., 2019; Lapin et al., 2019; Saile et al., 2020). Nevertheless, it is unknown how a signal is relayed from activated TNLs to RNLs *via* EDS1 complexes.

Aside from their beneficial function in conferring resistance to invading pathogens, NLRs may become activated erroneously, thus inducing autoimmunity (Bomblies et al., 2007; Chae et al., 2014). Autoimmune plants exhibit hallmarks of innate immune responses, such as marker gene induction and accumulation of the defense hormone salicylic acid. Additionally, plants are commonly dwarfed and exhibit curled leaves, tissue necrosis and spontaneous cell death (Bomblies et al., 2007; Alcazar and Parker, 2011; van Wersch et al., 2016). Autoimmunity has been described in at least three different scenarios: (*i*) transgenic lines ectopically expressing proteins (*e.g*. Xu et al., 2014; Stuttmann et al., 2016); (*ii*) mutant lines obtained in forward genetic screens (*e.g*. Li et al., 2001; Kim et al., 2010) and (*iii*) hybrids combining independently evolved alleles, then referred to as hybrid weakness or hybrid incompatibility (HI; Bomblies and Weigel, 2007).

In particular, intraspecific HI has been well-studied in Arabidopsis (*Arabidopsis thaliana*). Different degrees of HI, ranging from mild chlorosis to seedling lethality, occurred in approximately 2 % of all hybrids from systematic crossing of accessions representing most of the species’ diversity (Bomblies et al., 2007; Chae et al., 2014). Genetic dissection revealed that HI is generally induced by two genetically separable loci at least one of which encodes an NLR (Alcazar et al., 2009; Chae et al., 2014; Barragan et al., 2019). This pinpoints the plant innate immune system as a key player of HI. NLRs identified as causal in Arabidopsis HI include, but are not limited to, numerous *RECOGNITION OF PERONOSPORA PARASITICA* (*RPP*) loci. *RPP* loci were identified as conferring race-specific resistance to different isolates of the obligate biotrophic oomycete *Hyaloperonospora arabidopsidis (Hpa;* formerly *Peronospora parasitica*). For example, *RPP2* alleles present in reference accession Columbia (Col), but not those from accession Landsberg *erecta* (L*er*), confer resistance to *Hpa* isolate Cala2 (Sinapidou et al., 2004). Similarly, *RPP5* from accession L*er*, but not genes located at the homologous *RPP4/RPP5* locus in Col, confer resistance to *Hpa* isolate Noco2 (Parker et al., 1997; Noël et al., 1999). In line with the recognition-escape battle (“molecular arms race”) dictating natural host-pathogen systems, both *RPP* genes and the corresponding *Hpa* effectors occur in allelic series and are under strong selection (Botella et al., 1998; Rehmany et al., 2005). While gene clustering is rare in eukaryotes, *RPP* genes occur in eventually large and diversified clusters undergoing rapid evolution by conjunction of tandem duplications, illegitimate recombination and gene conversions (Jacob et al., 2013; van Wersch and Li, 2019), thus facilitating functional diversification towards new recognition specificities. Indeed, while NLR-coding genes anyways represent one of the most rapidly evolving gene families in plants, several *RPP* loci stand out as they were identified as particularly prone to variation and rearrangements in Arabidopsis (Van de Weyer et al., 2019; Jiao and Schneeberger, 2020; Lee and Chae, 2020). Altogether, diversified and rapidly evolving NLR loci thus appear particularly prone to induce autoimmunity in intraspecific HI.

The *RPP1* locus, also referred to as *DANGEROUS MIX2* (*DM2*), has been identified as a major hotspot for autoimmunity, and genetically interacts with at least seven distinct loci encoding NLR and non-NLR proteins (Alcazar et al., 2010; Tahir et al., 2013; Chae et al., 2014; Stuttmann et al., 2016). Interestingly, while *e.g*. RPP1^WsB^ (from accession Wassilewskija) and RPP1^Nd^ (from accession Niederzenz) confer *Hpa* resistance, beneficial immune functions were not reported for the autoimmunity-inducing “risk alleles” (originating from accessions L*er*, Uk-1, Bla-1, S. Tyrol and Dog 4). The *RPP1/DM2* locus in accession L*er* (*RPP1/DM2*^L*er*^) encodes seven to eight full-length TNLs. Respective genes are referred to as *RPP1-like R1-R8* or *DM2a-DM2h*, with the latter nomenclature utilized hereafter (Alcazar et al., 2009; Chae et al., 2014). *DM2*^L*er*^ causes HI when combined with alleles of the receptor kinase *STRUBBELIG-RECEPTOR FAMILY3* (*SRF3*) from the South Asian accessions Kashmir and Kondara (Alcazar et al., 2009; Alcazar et al., 2010). Furthermore, an EMS-induced allele of an O-acetylserine (thiol) lyase (*old3-1*) and ectopic expression of the immune regulator EDS1 in fusion with yellow fluorescent protein and a nuclear localization signal (EDS1-YFP^NLS^) induce autoimmunity in presence of *DM2^L*er*^* (Tahir et al., 2013; Stuttmann et al., 2016). We previously identified *dm2h* alleles in an EMS screen for genetic suppressors of EDS1-YFP^NLS^-induced autoimmunity and showed that *DM2h* was also essential for *SRF3*- and *old3-1*-induced autoimmunity (Stuttmann et al., 2016). Until recently, functional studies of risk loci were hampered by the complexity of the respective gene clusters, the similarity between the NLR genes and the strong dosage-dependent effects of transgenic expression of single NLRs. The contribution of risk loci to natural pathogen resistance and synergistic and/or epistatic interactions between risk locus-encoded NLRs remain therefore largely unexplored. In addition, the molecular mechanisms underlying autoimmune induction are unknown.

To explore the contribution of risk NLRs to pathogen resistance and autoimmunity and to reveal mechanisms of risk NLR activation, we genetically dissected the *DM2*^L*er*^ cluster, and re-constituted activation of a *DM2* risk NLR in heterologous expression assays. Relying on unique combinations of *DM2* genes generated by genome editing in the background of the native L*er* accession, we show that the single DM2h immune receptor is necessary and sufficient in all three cases of autoimmunity induced by EDS1-YFP^NLS^, *old3-1* and *SRF3*. However, we failed to uncover any relevance of the *DM2*^L*er*^-encoded TNLs for resistance to model pathogens. In addition, we functionally evaluated activation of a risk allele by an inducer, EDS1-YFP^NLS^, in *Nicotiana benthamiana* Agroinfiltration assays and in stable transgenic Arabidopsis. Our data suggest that activation of DM2h occurs by similar mechanisms as for classical sensor NLRs. Intriguingly, we provide evidence that the capacity to activate the NLR and to trigger autoimmunity is unrelated to the physiological activity of the inducer (here, EDS1/EDS1-YFP^NLS^). We therefore propose that autoimmune induction may, at least in some cases, rely on stochastic interactions of a non-native protein with an immune receptor. This emphasizes the importance of exercising caution when drawing conclusions regarding the immune function of a protein based on its ability to trigger an autoimmune response.

## Results

### Analysis of *nde-type* suppressor mutants: *DM2h* is the main locus required for autoimmunity induced by an EDS1-YFP^NLS^ fusion

Our screen for genetic suppressors of EDS1-YFP^NLS^-induced autoimmunity previously identified ^~^ 55 *<u>n</u>ear <u>d</u>eath <u>e</u>xperience* (*nde*) mutant lines (Stuttmann et al., 2016). To further elucidate the genetic requirements of autoimmune induction, we analyzed remaining *nde* lines. First, we Sanger-sequenced *DM2h* in 30 selected phenotypically stable *nde* mutant lines. Strikingly, 25/30 lines carried independent mutations within *DM2h* (Figure 1a, Table S1). We concluded that *DM2h* was the main gene required for EDS1-YFP^NLS^-induced autoimmunity targeted in our suppressor screen. The predicted *DM2h* gene model encompasses six exons, and encodes for a canonical TNL-type immune receptor (Figure 1a). TIR and NB-ARC domains are highly homologous among TNLs, and the three *nde* mutations mapping to these domains affected conserved residues (Figure S1). Most *nde* mutations affected residues located to the more divergent LRR domain and the LRR-adjacent C-terminal region (Figure 1a). Interestingly, we identified two nonsense mutations (W1129*, W1133*) that truncate DM2h by approximately 40 amino acids.

**Figure 1:**
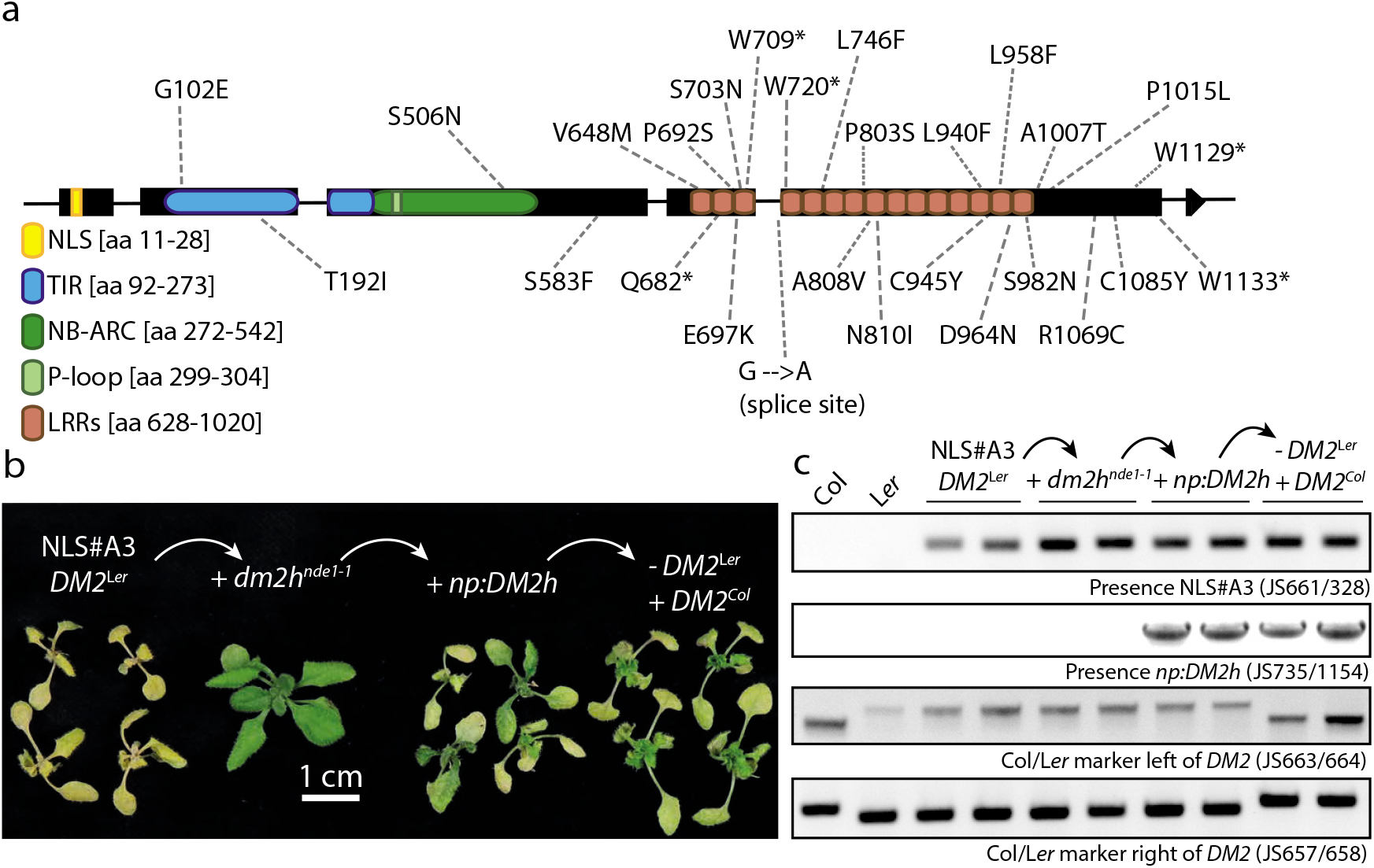
*DM2h* is the main locus targeted by the *nd*e suppressor screen, and is sufficient for EDS1-YFP^NLS^-induced autoimmunity. **a)** Protein motifs, domain boundaries and mutations discovered in different *nde* alleles parsed onto the *DM2h* gene model are shown (drawn to scale; gDM2h: 4031 nt ATG->STOP). A total of 15 LRRs is predicted for DM2h by LRRpredictor (Martin et al., 2020a). **b)** *DM2a-g*^L*er*^ are not required for autoimmunity induced by EDS1-YFP^NLS^. Plants were grown under high temperature conditions, and shifted to 18/16°C to induce autoimmunity for 8d prior to documentation. The NLS#A3 line (parental line used for the *nde* suppressors screen) expresses EDS1-YFP^NLS^ under *EDS1* promoter control in the Col *eds1-2* genetic background (a near isogenic line containing the *DM2*^L*er*^ region). The *dm2h^nde1-1^* allele was isolated from the suppressor screen, and was subsequently complemented by a T-DNA for expression of *gDM2h* under control of its own promoter (+ *np:DM2h*). From a cross to Col, a line containing both transgenes (EDS1-YFP^NLS^, *DM2h*), but not the *DM2*^L*er*^ region, was isolated. **c)** PCR-genotyping of lines used in b), showing presence/absence of the two transgenes (EDS1-YFP^NLS^, *np:DM2h*) and origin (Col, L*er*) of the *DM2* region.

From the remaining five *nde* lines harboring wild type *DM2h*, we focused on *nde2*. By a combination of mapping-by-sequencing (using SHORE; Schneeberger et al., 2009) and recombination mapping, we narrowed *nde2* down to an interval on chromosome 4 that lacked non-synonymous candidate SNPs according to whole-genome resequencing data. However, the candidate interval coincided with the insertion site of the EDS1-YFP^NLS^ transgene (At4g28490; Stuttmann et al., 2016). Thus, suppression of autoimmunity in *nde2* is most likely caused by downregulation of EDS1-YFP^NLS^ expression. Identification of causal mutations in remaining *nde* lines was not attempted.

Targeting of *DM2h* in most *nde* lines suggests that autoimmune induction by EDS1-YFP^NLS^ does not require additional *DM2*-encoded NLRs. We demonstrated this also by comparing autoimmunity in near-isogenic lines ectopically expressing *DM2h* and containing *DM2*^L*er*^ or *DM2*^Col^ (Figure 1b,c). A complementation line expressing *DM2h* under control of its native promoter in the *dm2h^nde1-1^* background (Col *eds1-2 dm2h^nde1-1^ np:gEDS1-YFP^NLS^ np:gDM2h*; Col *eds1-2* is a near isogenic line that also contains *DM2*^L*er*^; Stuttmann et al., 2016) was crossed to Col, and lines containing *DM2*^Col^ and both transgenes were selected (*EDS1-YFP^NLS^, DM2h;* Figure 1c). Autoimmunity was induced similarly when plants containing *DM2*^L*er*^ or *DM2*^Col^ (and ectopically expressing DM2h and EDS1-YFP^NLS^) were shifted to low temperatures (Figure 1b). Thus, the *DM2a-g* genes from the *DM2*^L*er*^ region have weak or no contribution to necrosis induction, and *DM2h* is necessary and sufficient for autoimmunity induced by EDS1-YFP^NLS^.

### Generation of *DM2*^L*er*^ mutant variants by genome editing

To dissect the functions of *DM2* genes in autoimmunity and resistance to pathogens, we generated derivatives of the *DM2*^L*er*^ cluster by genome editing (Ordon et al., 2017; Ordon et al., 2019). The genetic makeup of mutant lines is summarized in Figure 2 (see Figure S2 for details). *dm2c-4* and *dm2h-1* are single mutant lines. The *Δdm2a-g* mutant line expresses only *DM2h* from its native genomic locus. *Δdm2-11* (^~^70 kb deletion) and *Δdm2-3* (^~^120 kb deletion) lack all *DM2* genes, and also regions encompassing At3g44610-620 and At3g44680-700 were deleted in *Δdm2-3* (Figure 2). Some mutant lines were initially generated in the *old3-1* mutant background, and wild type *OLD3* (OASTL-A1; AT4G14880) was subsequently introduced by crossing to L*er* (Tahir et al., 2013; Ordon et al., 2017). Notably, *Δdm2-3* mutant plants (lacking *DM2* and flanking genes) were smaller at the seedling stage up to approximately 3-4 weeks and flowered earlier than control plants under short day conditions (Figure S2). These phenotypes were independent of *OLD3/old3-1*, and were not detected in *dm2-11* plants. Thus, the observed differences are likely due to deletion of the cluster-flanking genes in *Δdm2-3*.

**Figure 2:**
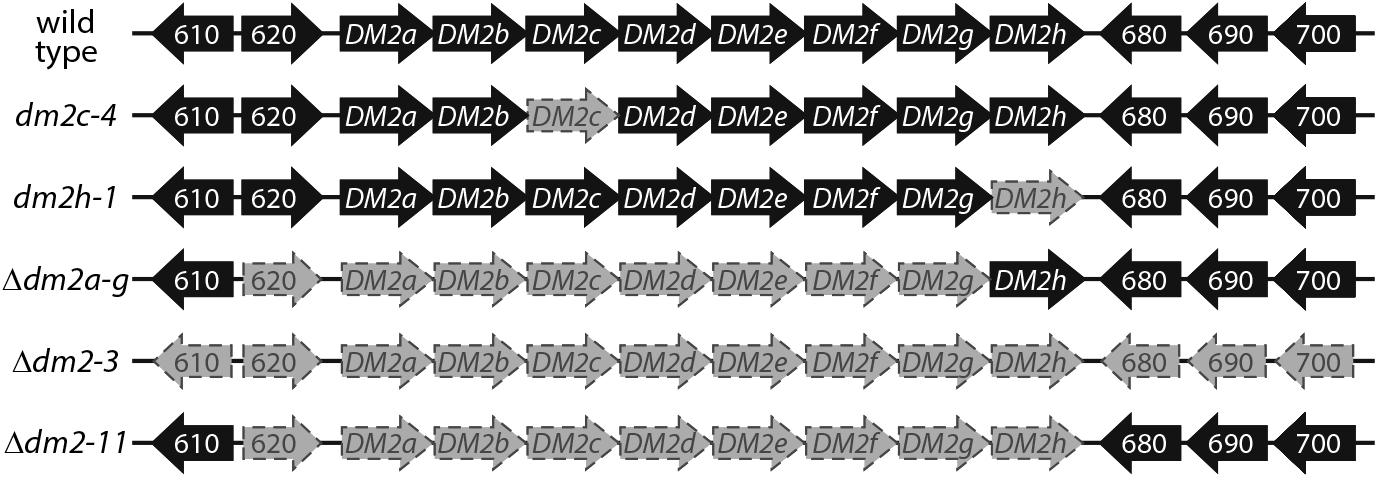
*dm2* mutant Arabidopsis lines used in this study. Schematic drawing of the *DM2*^L*er*^ region. Gene identifiers are the last three digits of At3g44xxx. Genes present in respective lines are depicted in black, and genes deleted or inactivated in mutant lines are shown in grey. Details on mutant lines are provided in Figure S2.

### The *DM2*^L*er*^ region contributes to oomycete resistance, but this is independent of the RPP1-like NLRs

Maintenance of the *DM2*^L*er*^ haplotype in natural populations despite its potentially detrimental effects on plant fitness in combination with certain genetic backgrounds (Alcazar et al., 2014; Atanasov et al., 2018) suggests it may confer selective advantages. We tested a potential role of the *DM2*^L*er*^ region in natural pathogen resistance by challenging Δ*dm2-3* mutant plants (lacking *DM2* and flanking genes) with virulent *Pseudomonas syringae* bacteria (*Pst* DC3000; Figure 3a) and compatible and incompatible *Hpa* isolates Cala2 and Emwa1, respectively (Figure 3b,c). The L*er rar1-13* line was included as a control for moderately impaired immunity (Muskett et al., 2002). As expected, *Pst* DC3000 bacteria grew better in *rar1-13* mutant plants, and also in *old3-1* mutant plants (compare Δ*dm2-3* and *Δdm2-3 old3-1;* Tahir et al., 2013). However, *Pst* DC3000 grew to similar levels in wild type and *Δdm2-3* mutant plants, indicating that the *DM2*^L*er*^ haplotype does not contribute to *Pst* DC3000 resistance (Figure 3a). In contrast, *Δdm2-3* mutant plants supported higher sporulation of the compatible *Hpa* isolate Cala2, and this phenotype was aggravated by presence of *old3-1* (Figure 3b). Similarly, isolate Emwa1, which is resisted by *RPP4/RPP8* in accession L*er* (Holub et al., 1994), showed enhanced outgrowth at infection sites and provoked less-confined, expanded HRs in *Δdm2-3* mutant plants in comparison to wild type L*er* (Figure 3c). This phenotype was more severe than that of the *rar1-13* mutant, which was only weakly impaired in resistance to Emwa1 under our conditions.

**Figure 3:**
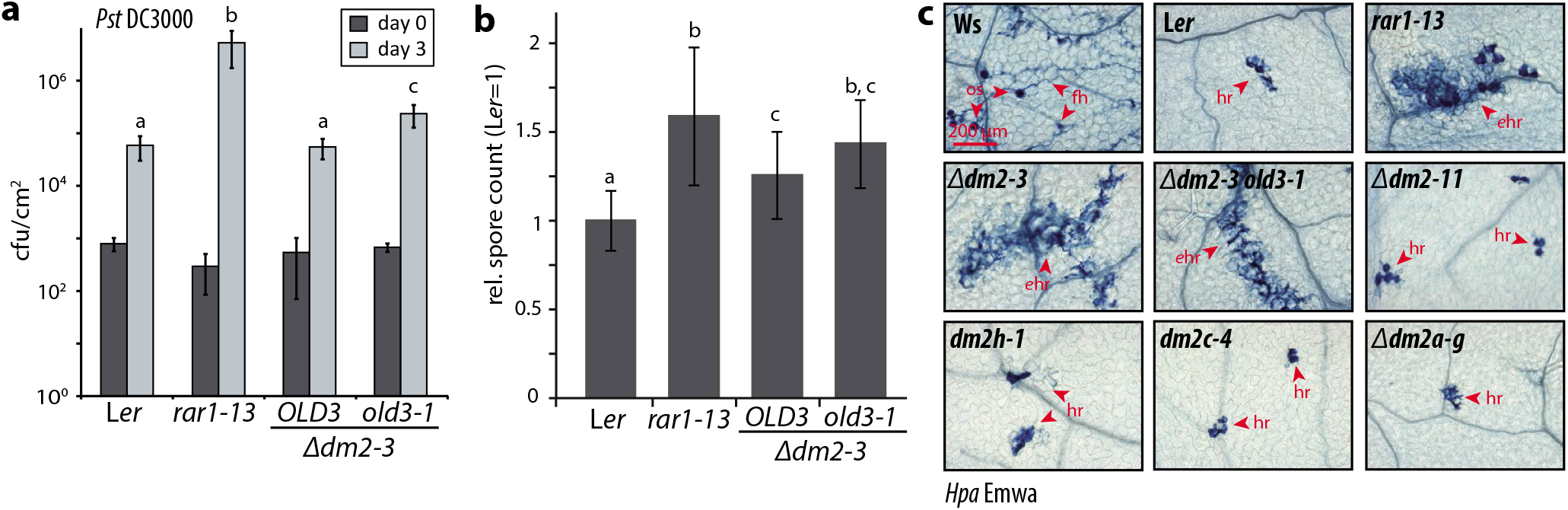
Infection of *dm2* mutant lines with bacterial and oomycete pathogens. **a)** Indicated plant lines were challenged with virulent *Pst* DC3000 bacteria, and bacterial titers determined at 0 and 3 dpi. Error bars indicate standard deviation of eight replicates, letters indicate statistically significant differences at day 3 as determined by one-way ANOVA and Fisher LSD (p < 0.001). The experiment was conducted three times with similar results; one representative experiment is shown. **b)** Indicated plant lines were challenged with virulent *Hpa* isolate Cala2. Sporulation was assessed 7 dpi to quantify pathogen growth. The experiment was conducted four times with four replicates. Data was normalized by arbitrarily setting sporulation on L*er* = 1, and all 16 replicates were included in the analysis. Error bars and statistics as in a). **c)** Infection phenotypes of indicated plant lines after challenge with *Hpa* isolate Emwa1, avirulent on L*er*. First true leaves were stained with Trypan Blue 6 dpi. The experiment was conducted five times. Representative micrographs are shown. os - oospores, fh - free hyphae, hr - hypersensitive response, ehr - expanded hr. Scale bar = 200 μm.

We assumed that one or several *Hpa* effectors might be weakly recognized by RPP1-like NLRs encoded at the *DM2*^L*er*^ locus. We used infection with *Hpa* isolate Emwa1 and Trypan Blue staining to analyze contributions of *DM2*^L*er*^ genes to *Hpa* resistance in more detail (Figure 3c). *Δdm2a-g, dm2c* and *dm2h* mutant plants were not impaired in resistance (Figure 3c). Unexpectedly, also the *Δdm2-11* mutant line, which carries a deletion of all *DM2* genes, was as resistant as L*er* wild type to *Hpa* Emwa1 (Figure 3c). This suggests that one of the flanking genes present in *Δdm2-11*, but absent in *Δdm2-3* (Figure 2), directly or indirectly contributes to *Hpa* resistance. *HISTONE DEACETYLASE9* (At3g44680) may be a most likely candidate, as it was reported to regulate flowering time (Figure S2; Kim et al., 2013; Park et al., 2019) and also plant resistance responses (Yang et al., 2020).

### DM2h is necessary and sufficient for autoimmunity induced by *old3-1* and *SRF3*^Kas/Kond^

The contribution of individual *DM2*^L*er*^ genes to temperature-dependent autoimmunity induced by *old3-1* or *SRF3*^Kas/Kond^ was not fully clarified (Alcazar et al., 2009; Alcazar et al., 2010; Tahir et al., 2013; Stuttmann et al., 2016; Atanasov et al., 2018). We therefore analyzed lines containing *old3-1* or *SRF3*^Kond^ and varying complements of *DM2* genes for autoimmune induction (Figure 4). Different lines were compared phenotypically, and salicylic acid (SA) levels or defense marker transcript levels were determined to quantitatively assess autoimmunity induction. There were no phenotypic differences between *old3-1* plants expressing only *DM2h* from its native genomic locus (*Δdm2a-g old3-1*) or those also containing additional *DM2* genes (Figure 4a,b). However, SA levels were mildly reduced in *Δdm2a-g old3-1* plants after temperature shift compared to *old3-1* plants (Figure 4b). Similarly, autoimmune hybrids obtained from crosses to Kondara (to introduce *SRF3*^Kond^) and containing either the full *DM2*^L*er*^ cluster or only *DM2h* (in the *Δdm2a-g* line) were phenotypically indistinguishable, and expression of the defense marker gene *Pathogenesis Related 1* (*PR1*) was slightly reduced in necrotic hybrids lacking *DM2a-g* (Figures 4c,d). In summary, the data do not suggest a major role for *DM2a-g*. Thus, *DM2h* is sufficient for autoimmune induction not only by EDS1-YFP^NLS^ (Figure 1), but also by *old3-1* and *SRF3*^Kond^.

**Figure 4:**
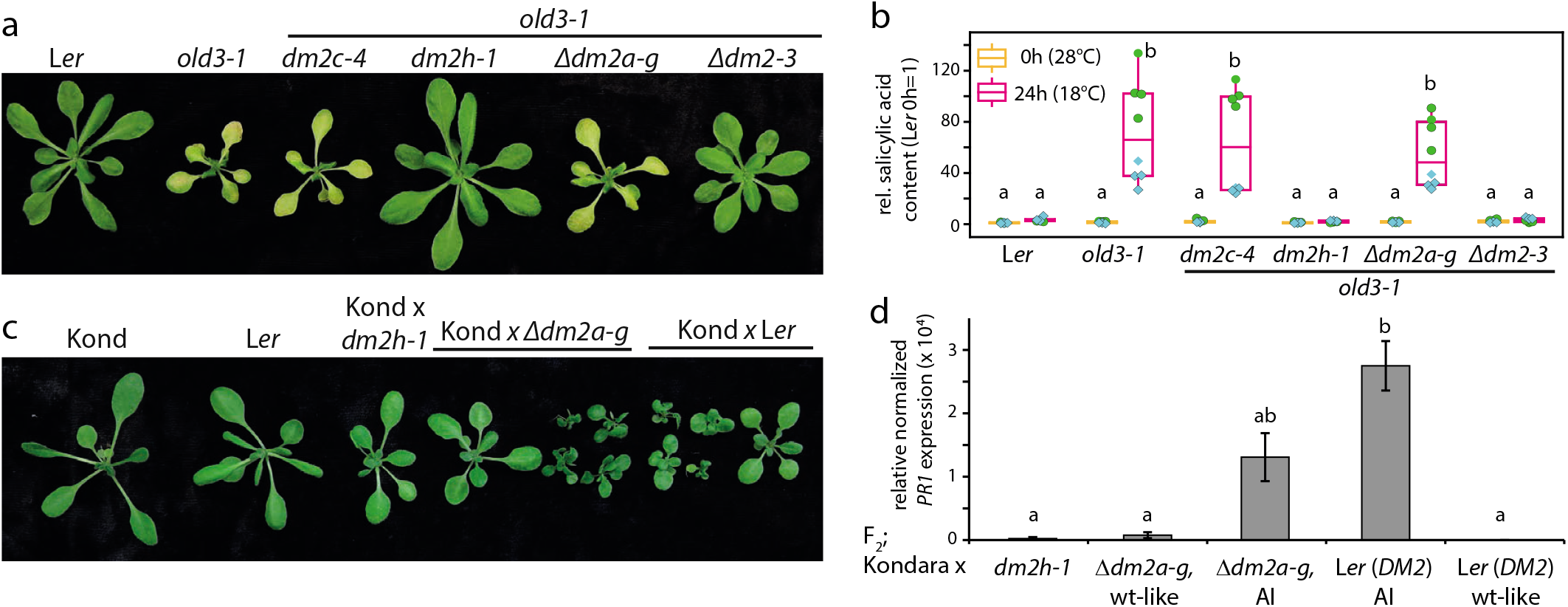
*DM2h* from the *DM2*^L*er*^ cluster is sufficient for autoimmune induction by *old3-1* or *SRF3*^Kond^. **a)** Contribution of *DM2* genes to *old3-1*-induced autoimmunity. Plants were grown for 7d under short day conditions, shifted for 14d to 28°C, and then shifted to 18/16°C. Phenotypes were documented 7d after temperature shift. **b)** Relative (L*er* 0h = 1) salicylic acid (SA) accumulation in leaf tissue of indicated plants lines at 28°C and 24 h after shift to 18/16°C. The experiment was conducted twice (green and cyan data points, respectively) with four replicates per experiment. Letters indicate statistically significant differences (ANOVA, Tukey *post-hoc* test, p<0,001). **c)** Contribution of *DM2* genes to *SRF3*^Kond^-induced autoimmunity. Control plants (Kond, L*er*) and F_2_ populations of the indicated crosses were grown under low temperature regime (14°C/12°C day/night; short day). Representative plants were documented after seven weeks, and both wild type-like and autoimmune plants are shown for segregating populations. Crosses were verified by PCR genotyping (Figure S4). **d)** Expression of the marker gene *PR1* in F_2_ plants from indicated crosses (to Kondara) as measured by quantitative RT-PCR. For crosses of Kond to L*er* and Δ*dm2a-g*, wt-like and autoimmune (AI) plants were analyzed. Means and standard errors of three biological replicates are shown. The experiment was conducted twice with similar results. Statistics as in b).

### Functional analysis of DM2h

Our genetic analyses showed that DM2h mediates autoimmunity induced by EDS1-YFP^NLS^, *old3-1* and *SRF3*^Kas/Kond^ (Figures 1,4). We wondered whether there were mechanistic differences between the risky DM2h NLR, prone to autoimmune induction, and previously characterized sensor TNLs. We therefore set out for functional analysis of DM2h.

We relied on heterologous expression in *Nicotiana benthamiana* (*Nb*; by Agroinfiltration) for functional analysis. On the one hand, we used a TIR domain fragment, DM2h_1-279_, which induces *EDS1*-dependent cell death in *Nb* (Gantner et al., 2019). On the other hand, we developed a co-expression assay to functionally interrogate the DM2h full length protein (Figure S4): Expression of DM2h together with EDS1-YFP^NLS^ or old3-1, but not expression of the proteins alone or co-expression of DM2h with EDS1-YFP or OLD3, induced HR-like cell death in *Nb*. Co-expression of DM2h with SRF3^Kond^, described to provoke autoimmunity recessively and only in absence of a compatible *SRF3* allele in Arabidopsis (Alcazar et al., 2010), did also not induce HR-like cell death. Thus, cell death induction in *Nb* faithfully recapitulated activation of DM2h-dependent autoimmunity in Arabidopsis, and can be used as a proxy for DM2h activation and initiation of TNL-mediated defenses.

*DM2h* encompasses an additional 5’exon present in some *RPP1* homologs (Figure 1a; Meyers et al., 2003; Chae et al., 2014). The encoded N-terminal extension contains a predicted myristoylation motif (including the critical G2 residue) and a bipartite nuclear localization signal (Stuttmann et al., 2016). Furthermore, DM2h contains a GK diresidue critical for ATP binding within the P-loop and a characteristic SH motif predicted to be required for TIR-TIR interactions (Bonardi et al., 2011; Williams et al., 2014; Burdett et al., 2019).

A variant of a DM2h_1-279_-GFP fusion carrying exchanges (to alanine) in the SH motif (SH-AA) accumulated to wild-type levels on western blots, but did not induce any plant reactions (Figure 5a,b), suggesting that DM2h cell death activity may involve TIR-TIR interactions. A DM2h_1-279_-GFP G2A variant was not stable (Figure 5b). G2S and G2E variants accumulated to low levels, but still induced residual cell death (Figure 5a,c), arguing against a requirement for myristoylation. This was further examined by co-expressing a DM2c^TIR^-DM2h^NB-LRR^ chimeric protein together with EDS1-YFP^NLS^ and old3-1 (Figure 5c). DM2c is highly similar to DM2h (^~^ 80 % amino acid identity in the region encoded by the DM2c/h chimeric construct), but lacks the N-terminal extension of DM2h and does not contain a predicted myristoylation motif or NLS. In co-expression assays, the DM2c^TIR^-DM2h^NB-LRR^ chimeric protein could mediate EDS1-YFP^NLS^- and old3-1-induced cell death, although this was mildly reduced in comparison to co-expression of native DM2h (Figure 5c). Taken together, these data suggest that the G2 residue is critical for stability, while myristoylation is not required for cell death executed by DM2h. Further, DM2h P-loop variants (G301A, K302R) were tested in co-expression assays (Figure 5d). While single amino acid exchanges strongly reduced the DM2h cell death activity, it was fully abolished when both residues were simultaneously exchanged in DM2h(GK-AR). DM2h, which often produced two signals on western blots, and P-loop variants accumulated to similar levels, suggesting that amino acid exchanges did not impair protein stability (Figure 5e; please note that cell death assays were conducted with untagged proteins, as tagged variants (*e.g*. N- and C-terminal 6xHA and GFP tags tested) were not or only weakly functional in co-expression assays).

**Figure 5:**
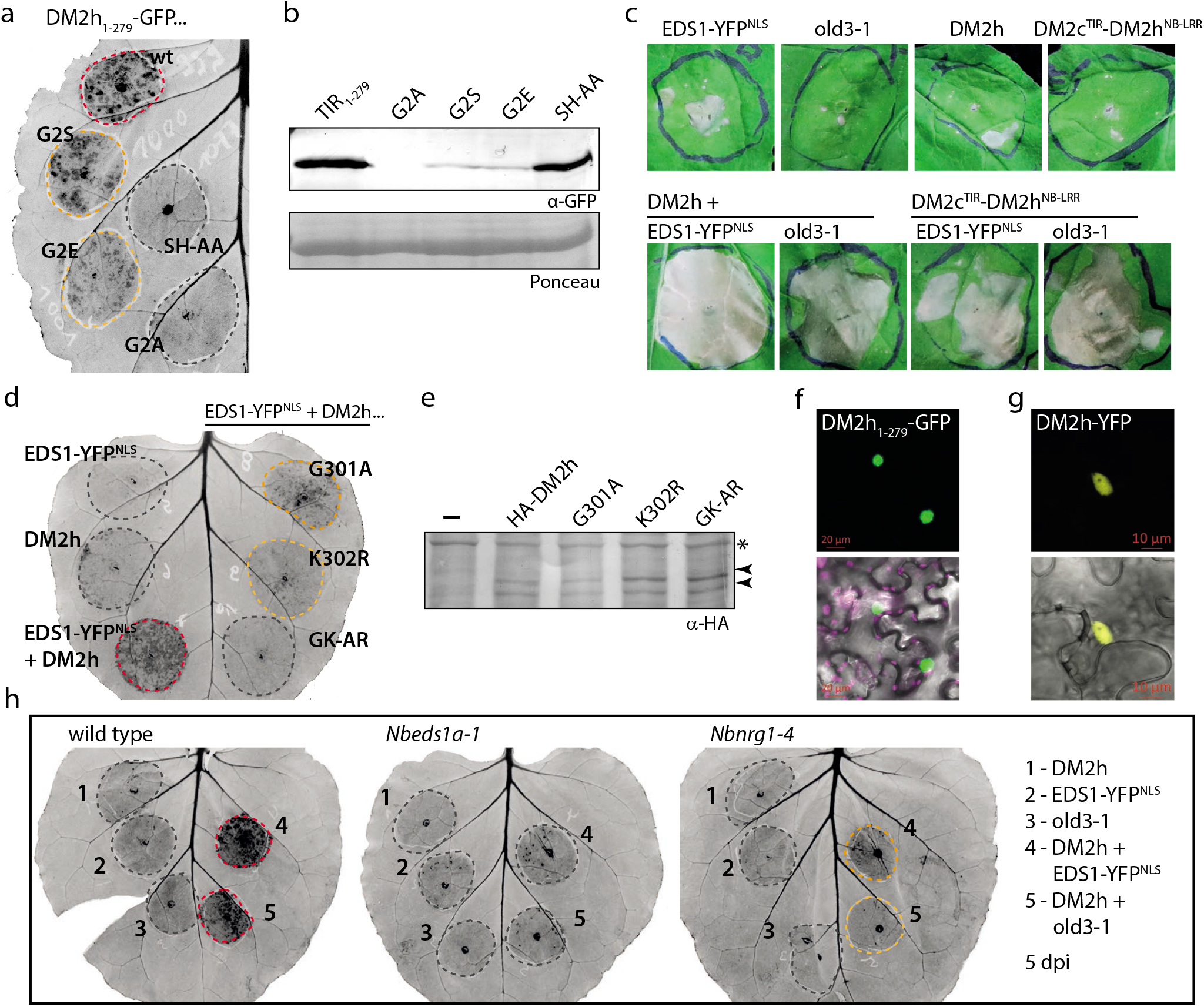
Functional analysis of the DM2h receptor. **a)** Cell death induction by DM2h_1-279_ and variants thereof in leaves of wild-type *N. benthamiana*. TIR domain fragments were expressed with a C-terminal GFP tag (35S:DM2h_1-279_-GFP), and plant reactions were documented 4 dpi. **b)** Immunodetection and accumulation of proteins transiently expressed in a). **c)** Cell death induction upon (co-) expression of DM2h or a DM2c/DM2h chimeric protein together with EDS1-YFP^NLS^ or old3-1. After Agroinfiltration, plants were incubated in the dark for 2d, plant reactions were documented 5 dpi. **d)** P-loop dependency of DM2h-mediated cell death. DM2h and P-loop variants, as indicated and without an epitope tag, were expressed by Agroinfiltration, and cell death induction was imaged 5 dpi. **e)** Immunodetection and accumulation of DM2h P-loop variants. Constructs similar to those used in d) but encoding an N-terminal 6xHA tag were used for Agroinfiltration. Protein extracts were prepared 3 dpi. Arrowheads mark two DM2h-specific signals. A non-specific signal, marked by an asterisk, is shown as loading control. **f)** Subcellular localization of the DM2h_1-279_ TIR domain fragment in *N. benthamiana*. GFP channel (upper panel) and a merged image (lower panel) including bright field and chlorophyll imaging (magenta) are shown. Similar results were obtained upon expression in *Nbeds1*. Scale bar = 20 μm. **g)** Subcellular localization of a DM2h-YFP fusion protein in transgenic Arabidopsis (Col *p35S:gDM2h-YFP*).14d old plants grown in short day conditions were used for imaging. With identical microscope settings, no fluorescence signal was detected in control plants. Multiple plants were analyzed; a representative micrograph is shown. Immunodetection of the DM2h-YFP fusion protein and growth phenotype of the transgenic line in Figure S5. **h)** DM2h-mediated cell death induction is dependent on the TNL downstream signaling components *EDS1* and *NRG1*. As in d), but different *N. benthamiana* mutant lines were used in co-expression assays. Details on the *nrg1-4* mutant line in Figure S6.

In addition, we aimed to determine the subcellular localization of DM2h by live cell imaging. No fluorescence signal was detected when the full length protein, in fusion to GFP, was expressed by Agroinfiltration. However, strong fluorescence in nuclei was observed when the DM2h_1-279_-GFP TIR fragment was expressed in *Nb* (Figure 5f). Similarly, DM2h was detected primarily in nuclei when examining a transgenic Arabidopsis line expressing DM2h-YFP under 35S promoter control in accession Col (Figure 5g).

Last, we determined the genetic requirements of DM2h-mediated cell death in *Nb* co-expression assays using an *Nbeds1* line and a newly generated *Nbnrg1* mutant line (Figure S6). Plant reactions induced upon co-expression of DM2h with EDS1-YFP^NLS^ or old3-1 were fully and partially abolished in *eds1* and *nrg1* mutant plants, respectively (Figure 5h).

Summarizing, our data suggest that induction of immune responses by DM2h may involve TIR-TIR interactions and requires an intact P-loop (Figure 5c,d), but not myristoylation (Figure 5a, c). Potential myristoylation is also not supported by localization studies, which suggest the nucleus as predominant compartment for DM2h (Figure 5f,g). Induction of downstream resistance responses by DM2h, here tested in *Nb*, requires EDS1 complexes and the helper NLR NRG1 (Figure 5h). Our data thus do not support mechanistic differences between the autoimmune risk NLR DM2h and classical sensor NLRs, as they recapitulate previously reported analyses of RPP1 orthologs functioning in *Hpa* resistance (Krasileva et al., 2010; Schreiber et al., 2016; Zhang et al., 2017) and also other sensor TNLs (e.g., Burch-Smith et al., 2007; Wirthmueller et al., 2007; Williams et al., 2014; Saile et al., 2020).

### EDS1-YFP^NLS^ operates upstream of DM2h

EDS1, in its function as an immune regulator, is assumed to signal at the level of or immediately downstream of activated TNL-type receptors. However, *Nb* co-expression assays (Figure 5) place *At*EDS1-YFP^NLS^ upstream of DM2h in cell death induction, as Arabidopsis EDS1 cannot exert immune signaling in *Nb* (Gantner et al., 2019; Lapin et al., 2019). This is also supported by the fact that EDS1-YFP^NLS^-induced cell death is abolished in *Nbeds1* plants (Figure 5h).

We made use of a previously identified loss-of-function allele of Arabidopsis EDS1, EDS1(F419E), to further disentangle EDS1 functions in downstream immune signaling from its role in activation of DM2h in the context of the EDS1-YFP^NLS^ fusion protein. EDS1(F419E) corresponds to a mutation identified in tomato EDS1 (*Sl*EDS1(F435E)), which abolishes EDS1 signaling functions, but does not interfere with assembly of EDS1 complexes with PAD4 or SAG101 (Gantner et al., 2019; Lapin et al., 2019). When expressed together with DM2h in *Nb*, EDS1-YFP^NLS^ and the corresponding F419E variant induced cell death to similar extents, while EDS1-YFP and EDS1-YFP^nls^ did not (Figure 6a). We also analyzed positioning of EDS1-YFP^NLS^ in Arabidopsis autoimmune induction by temperature shift experiments using primary transformants (Figure 6b). EDS1-YFP^NLS^ and an EDS1-YFP^NLS^(F419E, L258A, L262A) variant (under native promoter control) were combined with DM2h by transforming the constructs into wild type L*er* plants and/or the L*er eds1-2* mutant line. The L258A and L262A exchanges reduce the affinity of EDS1 to PAD4 and SAG101 (Wagner et al., 2013). They were combined with F419E in the transformation construct to avoid dominant negative effects from interference with the assembly of endogenous EDS1-PAD4/SAG101 complexes. EDS1-YFP^NLS^ induced autoimmunity among primary transformants in the L*er eds1-2* background, as expected (Figure 6b). In contrast, EDS1-YFP^NLS^(F419E, L258A, L262A) did not induce autoimmunity in *eds1-2* plants, but in wild-type L*er* background (Figure 6b). This confirms that EDS1(F419E) is not functional in downstream immune signaling (Lapin et al., 2019), and demonstrates it is still capable of activating DM2h to induce autoimmunity in presence of signaling-competent, endogenous EDS1 complexes in Arabidopsis. The F419E mutation did not interfere with protein stability or alter protein subcellular localization (Figure S7). Thus, cell death-based *Nb* assays and stable Arabidopsis transformants provide strong evidence for a position of EDS1-YFP^NLS^ upstream of DM2h in the activation of the immune receptor. Concomitantly, we conclude that DM2h activation by EDS1-YFP^NLS^ is unrelated to and independent of EDS1 immune signaling functions, albeit DM2h, as a classical TNL, requires EDS1 signaling functions for downstream signal relay.

**Figure 6:**
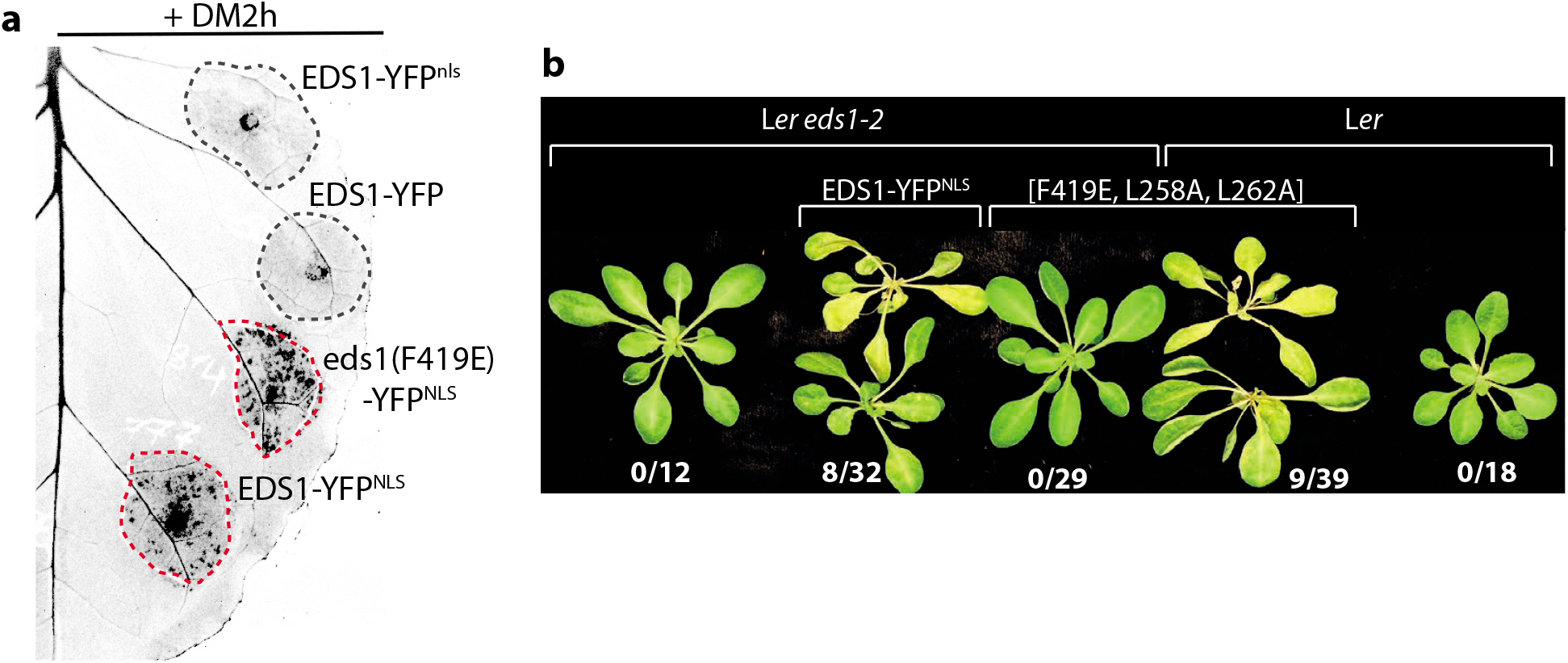
Activation of DM2h by EDS1-YFP^NLS^ is independent of EDSl’s role in signal transduction. **a)** Induction of HR-like cell death by different EDS1-YFP fusion protein variants in *N. benthamiana*. Indicated EDS1 variants were expressed together with DM2h, and cell death was imaged 6 dpi. Immunodetection of proteins and subcellular localization in Figure S7. **b)** Induction of autoimmunity by EDS1-YFP^NLS^ fusion protein variants. Transgenes for expression of immune-competent EDS1-YFP^NLS^ or the non-functional EDS1-YFP^NLS^(F419A, L258A,L262A) variant under control of the native promoter were transformed into accession L*er* or the L*er eds1-2* mutant line. Transgenic seeds were selected by FAST seed fluorescence. Plants were first grown four weeks under immune-suppressive conditions, and autoimmune induction was evaluated and documented 6d after shift to 18/16°C. Numbers indicate frequencies of autoimmune plants. Expression of transgenic proteins and presence/absence of endogenous EDS1 was shown on immunoblots (see Figure S7c).

### Identification of features within EDS1-YFP^NLS^ required for DM2h activation

The observations that DM2h functions similar to NLRs conferring natural pathogen resistance (Figure 5) and that EDS1-YFP^NLS^ operates upstream of the DM2h immune receptor (Figure 6) suggest that the fusion protein might mimic an effector activating a sensor NLR. We therefore hypothesized that EDS1-YFP^NLS^ contained a surface or epitope-like feature specifically recognized by the DM2h immune receptor.

We generated EDS1 expression constructs differing in the NLS (as EDS1-YFP did previously not induce autoimmunity; Figure 5; Stuttmann et al., 2016) or the linker connecting EDS1 with the YFP moiety (Figure 7a). Expression constructs encoded for an SV40 NLS lacking two terminal glycine residues, which were contained in the original EDS1-YFP^NLS^ fusion, or an NLS from c-myc, the human cancer protein. Additionally, constructs encoded either a 16 aa linker derived from the Gateway attB recombination site, or a short Ala-Ser-Ala (ASA) linker (Figure 7a). When tested in *Nb* co-expression assays, all fusion proteins containing the Gateway-linker activated DM2h and induced cell death, while those containing the ASA linker did not (Figure 7b). EDS1-YFP^NLS^ variants showed similar subcellular localizations, and were not impaired in protein accumulation (Figure S8). EDS1-YFP^NLS^ variants were further tested for their capacity to mediate TNL signaling and to induce autoimmunity in Arabidopsis. Constructs for expression of EDS1-YFP^NLS^ variants, under *EDS1* promoter control, were transformed into the Col *eds1-2* line (containing DM2^L*er*^, Figure 1b,c). Infection studies with primary transformants and *Hpa* isolate Cala2 showed that all variants could complement impaired *RPP2*-mediated resistance of the Col *eds1-2* line, and were thus functional in the context of downstream TNL signaling (Figure 7c). However, in agreement with results obtained in *Nb* co-expression assays, only those variants containing the Gateway linker also induced autoimmunity in temperature-shift experiments (Figure 7d). It is thus minor differences within the EDS1-YFP^NLS^ fusion protein that differentiate a DM2h-activating variant from one that is compatible with DM2h. These differences are not linked to EDS1 immune signaling functions.

**Figure 7:**
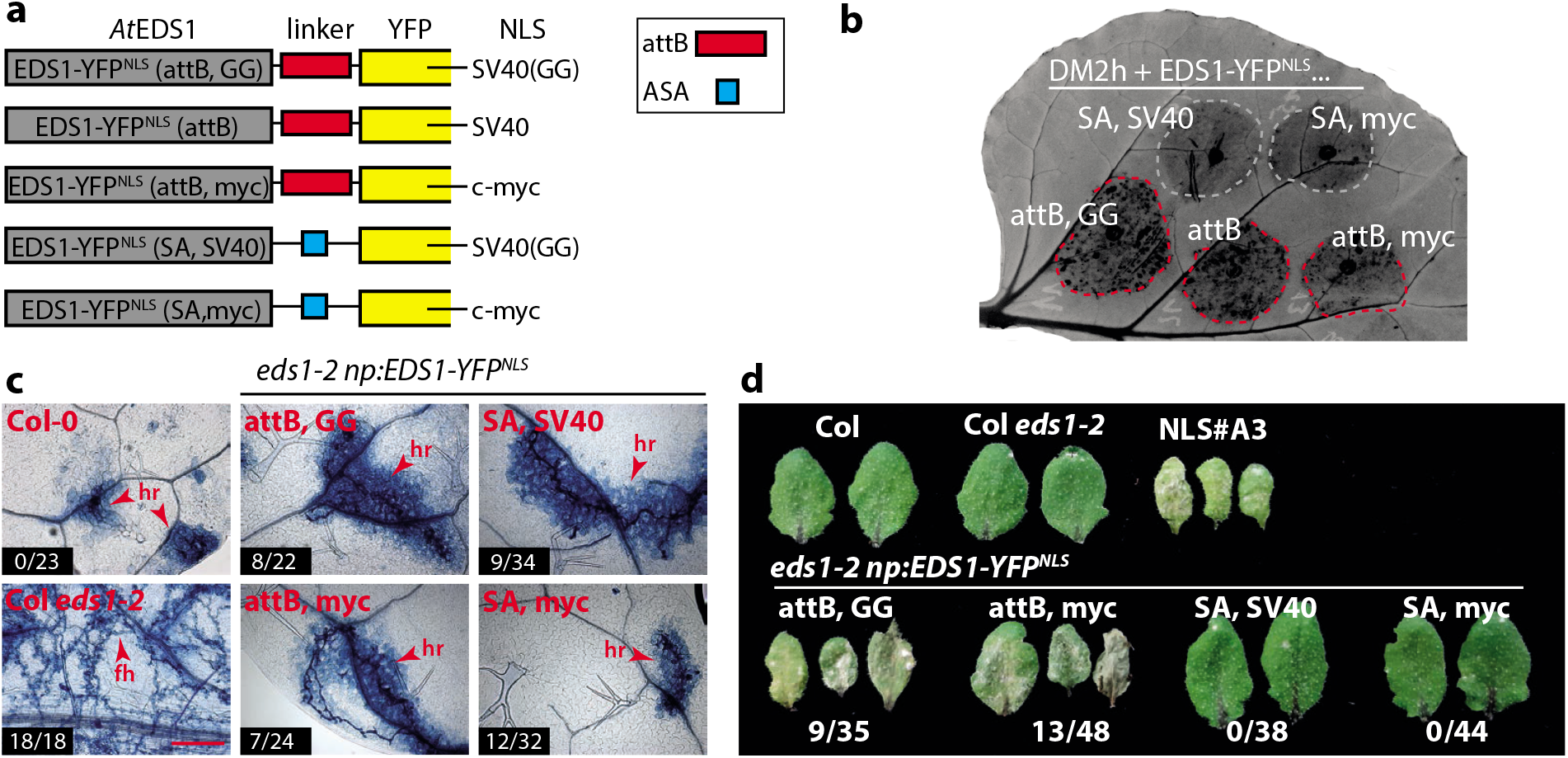
Analysis of EDS1-YFP^NLS^ properties required for activation of DM2h. **a)** Schematic representation of different EDS1-YFP^NLS^ variants compared for their capacity to activate DM2h. Red box represents the Gateway linker (KGGRADPAFLYKVVDG), blue box an Ala-Ser-Ala (ASA) linker. Precise sequence of used NLSs is as follows: SV40 - PKKKRKV*; SV40(GG) - PKKKRKVGG*; c-myc - SAPAAKRVKLD*. **b)** Induction of DM2h-mediated HR-like cell death by different EDS1-YFP^NLS^ variants in *N. benthamiana* co-expression assays. Plant reactions were imaged 5 dpi. Subcellular localization and accumulation of fusion proteins are shown in Figure S8. **c)** Functionality of EDS1-YFP^NLS^ variants in TNL signaling. Indicated variants were transformed, under control of the native *EDS1* promoter, into the Col *eds1-2* line. Primary transformants were selected by FAST seed fluorescence. Plants were grown for 7d under short day conditions and 10d under high temperature conditions prior to infection with *Hpa* Cala2. First true leaves were Trypan Blue-stained 6 dpi, and representative micrographs are shown. Numbers indicate plants with macroscopically visible sporulation. hr - hypersensitive response, fh - free hyphae. Scale bar = 200 μm. **d)** Autoimmune induction by EDS1-YFP^NLS^ variants. As in c), but plants were grown 7d under short day conditions, 20d under high temperature conditions and were then shifted to low temperature (18/16°C) for 5d, prior to documentation of autoimmune induction. Numbers indicate frequency of autoimmune plants among the total number of primary transformants that were analyzed. The experiment was performed twice. Frequencies for autoimmune plants were similar and total numbers from both replicates are indicated.

## Discussion

A number of NLR-coding loci, often referred to as risk loci or *DANGEROUS MIX* loci, are involved in induction of autoimmunity in diverse scenarios. Among these, DM2^L*er*^ is unique because of its role in at least three independent cases of autoimmunity, in genetic interactions with natural and EMS-induced alleles, or an ectopically expressed EDS1-YFP^NLS^ protein (Alcazar et al., 2009; Tahir et al., 2013; Stuttmann et al., 2016). In this study, the *DM2h* gene from the *DM2*^L*er*^ cluster was not only identified as the exclusive target of our EDS1-YFP^NLS^ suppressor screen (Figure 1), but also revealed as major driver of autoimmune induction when expressed from its endogenous genomic locus (in a *Δdm2a-g* mutant line) together with *old3-1* or *SRF3*^Kond^ (Figure 4; Atanasov et al., 2018). Thus, we demonstrate that a single NLR is necessary and sufficient in all three scenarios of autoimmune induction. This raises the questions how autoimmune risk NLRs may be different from classical sensor NLRs and how DM2h becomes activated in presence of different incompatible alleles.

### Sensor NLRs *vs*. risk-NLRs: Any evidence for conceptual differences?

RPP1-like NLRs encoded at the *DM2* locus in Arabidopsis accessions and conferring resistance to different *Hpa* isolates were extensively characterized in previous studies (Rehmany et al., 2005; Krasileva et al., 2010; Chou et al., 2011; Steinbrenner et al., 2015; Goritschnig et al., 2016; Schreiber et al., 2016; Zhang et al., 2017; Horsefield et al., 2019; Wan et al., 2019). Summarizing our current understanding, activation of RPP1-dependent immune responses depends on *i*) direct interaction of the recognized effector with the receptor’s LRR domain, *ii*) receptor oligomerization, controlled by ATP-binding at the P-loop, *iii*) TIR-TIR interactions *via* the AE and DE-interfaces, *iv*) enzymatic (NADase) activity dependent on a critical TIR domain glutamate residue. It is tempting to speculate that also for RPP1, formation of resistosome complexes regulates the TIR domain enzymatic activity (Martin et al., 2020b). Furthermore, RPP1-mediated resistances depend on EDS1 complexes and helper NLRs (Wagner et al., 2013; Qi et al., 2018; Jubic et al., 2019). Here, we reinforce and add to previous analyses of the RPP1-like risk NLRs DM2d^Uk-1^ and DM2h^Bla-1^ (Chae et al., 2014; Tran et al., 2017) by functional analysis of DM2h^L*er*^ (Figure 5). Our results support that also DM2h activation depends on a functional P-loop and TIR-TIR interactions. Additionally, our *Nb* assays reveal that DM2h-mediated responses are, fully and partially, dependent on presence of EDS1 complexes and the NRG1 helper NLR, respectively. Last, we identify the nucleus as most likely compartment for DM2h localization (Figure 5h,g), which is in agreement with previous analyses of the TNLs N and RPS4 (Burch-Smith et al., 2007; Wirthmueller et al., 2007). Overall, our results do thus not support differences, at the mechanistic level, between autoimmune risk-NLRs and sensor NLRs characterized in pathogen resistance.

In absence of mechanistic differences, what could be the basis for the frequent involvement of certain NLR loci, such as *DM2*, in autoimmunity? The *DM2* locus and additional complex *NLR* loci stand out, at the genome-scale, for their complexity and evolutionary dynamics (Chae et al., 2014; Van de Weyer et al., 2019; Jiao and Schneeberger, 2020; Lee and Chae, 2020). Thus, a disproportionally large number of diversified RPP1-like NLRs may be encoded within the global Arabidopsis gene pool, which may explain to some extent the incidence of *DM2* and other complex *NLR* clusters in autoimmunity. An alternative explanation for why certain highly variable NLRs are particularly prone to autoimmune induction could be that they differ in their activation kinetics. In the equilibrium-based switch model, NLRs cycle between the ADP-bound OFF and the ATP-bound ON state also in absence of a recognized effector (Bernoux et al., 2016). However, baseline activation levels differ between NLRs, and signaling capacities can be tuned to less responsive or hyper-responsive by altering intramolecular interactions (Qi et al., 2012; Bernoux et al., 2016). Risk NLRs may, at least in some cases, be shifted particularly far towards the ON/ hyper-responsive state in comparison to evolved sensor NLRs. This may place them directly at the brink of full activation, so that spurious interactions or even cellular perturbations may be sufficient for immune receptor activation and resistosome formation. The concept of cellular perturbation as an NLR trigger is, *e.g*., supported by an accession-specific *TNL* allele inducing autoimmunity under osmotic stress (Ariga et al., 2017). Activation of NLRs by altered cell physiology could provide one possible explanation for the apparent promiscuous activation properties of DM2h as well as other risk NLRs, such as SUPPRESSOR OF NPR1-1, CONSTITUTIVE1 (SNC1) (see *e.g*. Chakraborty et al., 2018).

### DM2h as a guardian NLR, or direct binder?

The most plausible scenarios for the induction of DM2h-mediated autoimmunity are direct activation of the DM2h immune receptor through physical interaction with gene products of incompatible alleles or disturbance of guard-guardee pairs (Rodriguez et al., 2016; Kourelis and van der Hoorn, 2018). While experimental data supporting either scenario are scarce, we think that a number of arguments favor direct activation. RPP1 receptors conferring *Hpa* resistance directly bind corresponding ARABIDOPSIS THALIANA RECOGNIZED1 (ATR1) effector proteins *via* their LRR domain, and variation at *RPP1* and *ATR1* loci and structure-function analyses support ongoing co-evolution of receptor and ligand surfaces (Rehmany et al., 2005; Krasileva et al., 2010; Chou et al., 2011; Steinbrenner et al., 2015; Goritschnig et al., 2016). There does not appear to be any difference *per se* between NLRs encoded at risk loci and those with characterized immune functions, and inducers of autoimmunity demonstrate the same behavior of allele-specific receptor activation as pathogenic ATR1 effectors (Figures 5,7; Tahir et al., 2013; Alcazar et al., 2014; Chae et al., 2014). Genetic screens have so far also not revealed further components required for DM2h-mediated autoimmunity, as might be expected for a scenario of indirect activation (this work; Stuttmann et al., 2016; Atanasov et al., 2018). Maybe most significantly, there is little support for a function of DM2h as guardian NLR from an evolutionary perspective. Guardian NLRs can be expected to co-evolve with guardees and thus to experience balancing or purifying selection. In contrast, NLRs directly binding cognate effectors are required to rapidly evolve novel specificities to adapt to newly arising effector alleles, and thus underlie diversifying selection – which is clearly the case for RPP1-like NLRs. Indeed, Prigozhin et al. (2020) recently proposed that refined NLR clade assignment, coupled with analysis of amino acid diversity in near-allelic series, can be used to computationally discriminate guardian NLRs and direct binders (Prigozhin and Krasileva, 2020). In this analysis, a class of highly variable NLRs contained all the described Arabidopsis autoimmune loci (including *DM2h*), but not a single guard NLR (Prigozhin and Krasileva, 2020).

It is also worth noting that two of the EMS-induced *dm2h* alleles identified in the *nde* mutants, E697K and S703N (Figure 1), locate to the presumably surface-exposed positions on the concave surface of the DM2h-LRR domain (variable positions within the LxxLxLxx LRR consensus sequence, prediction of DM2h LRRs using NLRpredictor; Martin et al., 2020a). These mutations may highlight an interface for the binding of EDS1-YFP^NLS^ on the DM2h-LRR, and it will be interesting to test whether the respective alleles might be specific for EDS1-YFP^NLS^ recognition; in other words, whether they maintain responsiveness to old3-1 and/or *SRF3*^Kond^. Furthermore, a total of six independent *dm2h* alleles (including two minor truncations) also highlight the importance of the C-terminal non-LRR domain for autoimmune induction by EDS1-YFP^NLS^. In the recently solved Roq1 cryo-EM structure, this Post-LRR domain contributes to ligand binding together with the LRR (Martin et al., 2020b).

We could not find any support for direct interaction of old3-1 and EDS1-YFP^NLS^ with DM2h or domains thereof in initial yeast two hybrid and co-immunoprecipitation assays (not shown). However, more sensitive interaction assays suitable for transient interactions, such as split Luciferase or proximity labeling by TurboID (Luker et al., 2004; Branon et al., 2018; Mair et al., 2019; Arora et al., 2020), may be required for revealing a direct interaction between DM2h and inducers, as it is known that also NLR-effector interactions in pathogen recognition are often highly transient and difficult to capture experimentally (Saur et al., 2019; Song et al., 2020).

### Role of autoimmunity risk loci in natural pathogen resistance: Adaptive benefits?

Despite the potential risk of autoimmunity posed by the highly variable NLRs, they are maintained in the natural gene pool (Bomblies and Weigel, 2007; Alcazar et al., 2014). This suggests they may provide selective advantages in a given environment, potentially also due to a low activation threshold. Here, using CRISPR/Cas lines with different complements of *DM2* genes, we failed to show any role of this locus in pathogen resistance (Figure 3). This is reminiscent of a previous analysis of *DM2* haplotypes (Atanasov et al., 2018). However, in addition to evolutionary and population genetics analyses supporting functional relevance of risk loci (*e.g*. Chae et al., 2014), amino acid diversity within the LRRs of highly variable NLRs supports that variation does not occur randomly, but is the result of underlying selective pressure. For example, amino acid diversity is higher at surface-exposed positions within the LRR fold and often involves hydrophobic residues which may mediate protein-protein interactions (Prigozhin and Krasileva, 2020). Lastly, considering that risk loci are commonly selected for in breeding programs (Bomblies and Weigel, 2007), we most likely lack appropriate pathosystems to detect beneficial functions of risk loci, or effects may be subtle or masked by redundancy in incompatible interactions. To that end, recent system analyses suggest that 68 % of plant pathogenic *Pseudomonas* strains are resisted by multiple NLRs in Arabidopsis (Laflamme et al., 2020).

### Activation of risk-NLRs by incompatible alleles: EDS1-YFP^NLS^ as a case study

Our data show that, although EDS1 is a well-known immune regulator required for signaling downstream of activated TNLs, the EDS1-YFP^NLS^ fusion protein operates upstream of the DM2h immune receptor to induce autoimmunity (Figures 5,6). DM2h activation depends on very specific properties of the fusion protein, *i.e*. the presence of the Gateway-linker and an NLS, but is independent of EDS1 immune signaling functions (Figures 6,7). Theoretically, DM2h activation could result from disturbance of a guard-guardee pair - EDS1-NLR interactions were reported and also guarding of EDS1 proposed (*e.g*. Bhattacharjee et al., 2011; Huh et al., 2017) – but, in particular, heterologous *Nb* assays argue otherwise. Case in point, DM2h activation can be reconstituted in *Nb*, although EDS1 proteins and functions have diverged between Arabidopsis and *Solanaceae* (Gantner et al., 2018; Lapin et al., 2019). Accordingly, induction of autoimmunity, at least in this case, is not predictive of immune functions of an incompatible allele. Similarly, activation of DM2h by old3-1 appears to depend on very specific properties of the incompatible allele (a point mutation); an *old3* loss-of-function T-DNA allele does not induce autoimmunity (Tahir et al., 2013). Notably, also the activation of immune receptors involved in dominant HI can often be reconstituted in heterologous *Nb* assays, and is limited to very specific alleles of the non-NLR locus in populations (*e.g*. Chae et al., 2014; Tran et al., 2017; Li et al., 2020). We therefore propose that autoimmunity in negative epistatic interactions involving NLRs may, at least in some cases, occur stochastically due to the recognition of corresponding patterns by NLRs. This shall function as a cautionary note for interpretation of dominant autoimmunity, in particular upon involvement of highly variable and thus risky NLRs.

## Material and Methods

### Plant material and growth conditions

*N. benthamiana* plants were cultivated in a greenhouse with a 16-h light period (sunlight and/or IP65 lamps (Philips) equipped with Agro 400 W bulbs (SON-T); 130–150 μE/m^2^*s; switchpoint; 100 μE/m^2^*s), 60% relative humidity at 24/20°C (day/night). The *Nbeds1 (Nbeds1a-1*) mutant line is published (Ordon et al., 2017). Details on *Nbnrg1* mutant lines are provided in Figure S3. *Arabidopsis thaliana* plants were cultivated under short day conditions (8h light, 23/21°C day/night, 60 % relative humidity) or in a greenhouse under long day conditions (16h light) for seed set. For suppression of autoimmunity, plants were germinated under short day conditions (7-8 d) and subsequently cultivated in a growth chamber with 28/26°C day/night temperatures and 12h light (^~^ 120 μE/m^2^*s). For temperature-shift experiments, plants were moved into a growth chamber with 18/16°C day/night cycle (12h illumination). For induction of hybrid necrosis (*SRF3*-*DM2*^L*er*^), plants were grown with 14/12°C day/night temperatures under short day conditions for 4 – 6 weeks. Arabidopsis mutant or transgenic lines in the Columbia background (Col *eds1-2*, EDS1-YFP^NLS^#A3 (in *eds1-2), dm2h^nde1-1^* (in EDS1-YFP^NLS^#A3), *np:DM2h* (in *dm2h^nde1-1^*) were previously described (Bartsch et al., 2006; Stuttmann et al., 2016). Mutant lines in the Landsberg *erecta* (L*er*) background used in this study were *eds1-2* (Aarts et al., 1998), *rar1-13* (Muskett et al., 2002) and several lines deficient in genes of the *DM2* cluster. *DM2*-deficient lines were reported previously (Ordon et al., 2017; Ordon et al., 2019) or details are provided in Figure S2. Furthermore, Arabidopsis accessions Kashmir and Kondara were used.

### Sequencing of *nde1* alleles, *DM2h* cDNA cloning and multiple sequence alignments

*DM2h* was PCR-amplified in two fragments using oligonucleotides JS724/1156 and JS1155/725 (Table S2). Amplicons were purified using a column-based kit and sequenced using oligonucleotides JS1153/729/731 and JS733/1381, respectively. To clone the *DM2h* cDNA, *DM2h* was first transiently expressed in *Nb* using a gDNA fragment under 35S promoter control. Tissues were used for RNA extraction 2 dpi, and cDNA was generated using RevertAid Reverse Transcriptase (Thermo). *DM2h* was amplified from cDNA in five fragments to eliminate internal *BpiI* restrictions sites (domestication) during cloning, and also to prune the second intron, which was retained in several fragments cloned from cDNA prepared from *Nb* (Primers JS972/973, JS729/1035, JS1036/974, JS975/976, JS977/978). Fragments were subcloned in pUC57 (cut/ligation with either *SmaI* or *Eco*RV), sequence-verified and combined in pICH41308 by a *Bpi*I cut/ligation reaction (Engler et al., 2014). The final construct, pJOG273, was again verified by sequencing, and used as template for all further clonings. Multiple sequence alignments were generated using tcoffee (http://www.tcoffee.org/), and rendered using ESPript (http://espript.ibcp.fr; Robert and Gouet, 2014).

### Infection assays and salicylic acid measurements

Infection assays with *Hyaloperonospora arabidopsidis* (isolates Emwa and Cala) were conducted as previously described (Stuttmann et al., 2011). For infection assays with *Pst* DC3000, bacteria resuspended in 10 mM MgCl_2_ were syringe-infiltrated into leaves of 4 – 6 week-old Arabidopsis plants at an OD_600_ = 0.0002. For day 0, samples were taken ^~^ 3h after infiltration, and 4 replicates each consisting of 3 leaf discs were processed. On day 3, 8 replicates were used. Leaf discs were shaken in 10 mM MgCl_2_ containing 0.01 % Silwet for 2 – 3h, serial dilution series were prepared and plated for determination of colony forming units (cfu). For statistical analyses, data was normalized and/or transformed, and ANOVA with Tukey post-hoc test were used. Metabolite extractions for SA measurements were done essentially as described previously (Balcke et al., 2017). Briefly, homogenized frozen material was cryo-extracted twice using 900 μL cold dichloromethane/ethanol and 150 μL aqueous hydrochloric acid buffer (pH 1.5). The organic phase was collected and the extraction residue was extracted again with tetrahydrofurane. Both extracts were combined and dried in a nitrogen stream. The dried residues were resuspended in 180 μL 80% methanol and centrifuged before injection. SA was measured using a Nucleoshell RP18 column (Macherey & Nagel) on a Waters ACQUITY UPLC System (flow rate 400 μL/min, column temperature 40 °C). Eluents A and B were aqueous 0.3 mmol/L NH4HCOO (adjusted to pH 3.5 with formic acid) and acetonitrile, respectively. Elution was performed as following: 0-2 min 5% eluent B, 2-19 min linear gradient to 95% B, 19-24 min 95% B. Mass spectrometric detection after electrospray ionization was performed *via* MS-TOF-SWATH-MS/MS (TripleToF 5600, both AB Sciex GmbH) operating in negative ion mode and controlled by Analyst 1.6 TF software. Source operation parameters were as following: ion spray voltage, −4500 V; nebulizing gas, 60 psi; source temperature, 600 °C; drying gas, 70 psi; curtain gas, 35 psi. SA was quantified using PeakView based on the [M-H] precursor m/z of 137.024 ± 0.02 Da and qualified *via* an authentic standard.

### Transient protein expression, immunodetection and live cell imaging

Plate-grown bacteria (Agrobacterium strain GV3101 pMP90 or pMP90RK) were resuspended in Agro Infiltration Medium (AIM; 10 mM MES pH 5.8, 10 mM MgCl_2_) and infiltrated with a needless syringe at OD_600_ = 0.6 per strain. Cell death-based assays in *Nb* were repeated at least five times for evaluation of phenotypes, and detection of expressed fusion proteins was conducted for at least two replicates. For UV light documentation, *Nb* leaves were photographed on a gel documentation system (AlphaImager®) using 5s exposure time (aperture 2.0). Image adjustments were applied to full images. Trypan Blue staining was done as previously described (Stuttmann et al., 2011). For incubation of *Nb* plants in the dark after infiltration, whole plants were placed in a growth cabinet (24/20°C, 16h/8h, 60 % RH) without illumination for 2d, and cell death was documented 5 dpi. Samples for immunodetection of fusion proteins were taken 3 dpi. Leaf discs were ground in liquid nitrogen, leaf powder resuspended in Laemmli buffer and proteins denatured by boiling. Total protein extracts were separated by SDS-PAGE and blotted on nitrocellulose membranes. Primary antibodies were α-GFP (mouse) and α -HA (rat; both from Roche), α –EDS1 (Rabbit) and horseradish peroxidase-coupled (GE Healthcare) or alkaline phosphatase-coupled (Sigma-Aldrich) secondary antibodies were used. A Zeiss LSM780 confocal laser scanning microscope was used for live cell imaging. All images are single planes.

### Plant transformation and genome editing

Arabidopsis was transformed by floral dipping as previously described (Logemann et al., 2006), and primary transformants were either selected by resistance to phosphinotricin (BASTA) or seed fluorescence (FAST; Shimada et al., 2010). *Nb* was transformed as previously described (dx.doi.org/10.17504/protocols.io.; Gantner et al., 2019), using previously described target sites / sgRNAs for *NRG1* editing (Qi et al., 2018). Different versions of pDGE vectors (Ordon et al., 2017; Ordon et al., 2019; Barthel et al., 2020) were used to generate *dm2* and *Nbnrg1* mutant lines by *Sp*Cas9. Respective target sites are provided in Figures S2 (Arabidopsis lines) and S3 (*Nbnrg1*). sgRNAs were prepared by cloning of hybridized oligonucleotides in pDGE shuttle vectors, as previously described (Ordon et al., 2017). Mutant lines were cured of the editing construct, with the exception of the *dm2-11* line.

### Molecular cloning, genotyping and quantitative RT-PCR

The GoldenGate strategy (Engler et al., 2008) was used for most clonings. Different DNA modules from the Modular Cloning system, the Plant Parts I and the Plant Parts II collections were used, and cut/ligation reactions employing either *BsaI* or *BpiI* together with T4 DNA Ligase (Thermo, 1-5 u/μl) were conducted using approximately 20 fmol of each DNA module as previously described (Weber et al., 2011; Engler et al., 2014; Gantner et al., 2018). Additional plasmids were generated using Gateway cloning. In these cases, entry clones were generated by PCR (via TOPO cloning or GoldenGate cloning), and inserts were subsequently recombined into expression plasmids by LR reactions. Detailed information on plasmids used and/or generated in this study are provided in Table S3. A lab-internal Taq polymerase preparation was used for genotyping of different Arabidopsis lines. Genomic DNA was extracted by a CTAB protocol, and primer sequences are provided in Table S2. For transcript analyses, RNA was extracted by a TRIzol protocol, as previously described (Gantner et al., 2019). The Reverse Transcriptase Core Kit was used for cDNA synthesis, and the Takyon No ROX SYBR 2X MasterMix Blue dTTP qPCR Kit (both Eurogentech) for quantitative real-time PCR using a CFX96 detection system (Bio-Rad). Transcript data was normalized to *UBQ10*; primer sequences are provided in Table S2.

## Supporting Information

**Table S1:** Molecular details on identified *dm2h/nde1* alleles

| mutant allele | mutation | phenotype |
| --- | --- | --- |
| *nde1-1* | Δ16bp | wild type-like (Stuttmann et al., 2016) |
| *nde1-3* | Δ*dm2c-h* | wild type-like (Stuttmann et al., 2016) |
| *nde1-13* | W1129STOP | wild type-like (Stuttmann et al., 2016) |
| *nde1-150* | R1069C | wild type-like (Stuttmann et al., 2016) |
| *nde1-175* | C945Y | wild type-like (Stuttmann et al., 2016) |
| *nde1-201* | A1007T | n. d. |
| *nde1-203* | L940F | n. d. |
| *nde1-8* | L958F | n. d. |
| *nde1-18* | P692S | partial rescue (intermediate) |
| *nde1-20* | P1015L | n. d. |
| *nde1-23* | P803S | wild type-like |
| *nde1-24* | G102E | wild type-like |
| *nde1-26* | L746F | n. d. |
| *nde1-31* | recombination event at *DM2* | n. d. |
| *nde1-32* | V648M | wild type-like |
| *nde1-34* | S506N | partial rescue (intermediate) |
| *nde1-40* | S703N | wild type-like |
| *nde1-41* | W709STOP | wild type-like |
| *nde1-49* | W720STOP | partial rescue (intermediate) |
| *nde1-51* | splice site | weak |
| *nde1-53* | D964N | wild type-like |
| *nde1-59* | A808V | wild type-like |
| *nde1-60* | W1133STOP | wild type-like |
| *nde1-65* | Q682STOP | wild type-like |
| *nde1-75* | C1085Y | wild type-like |
| *nde1-114* | S583F | n. d. |
| *nde1-145* | E697K | wild type-like |
| *nde1-147* | S982N | wild type-like |
| *nde1-161* | T192I | wild type-like |
| *nde1-166* | N810I | wild type-like |

n.d. – not determined

**Table S2:**
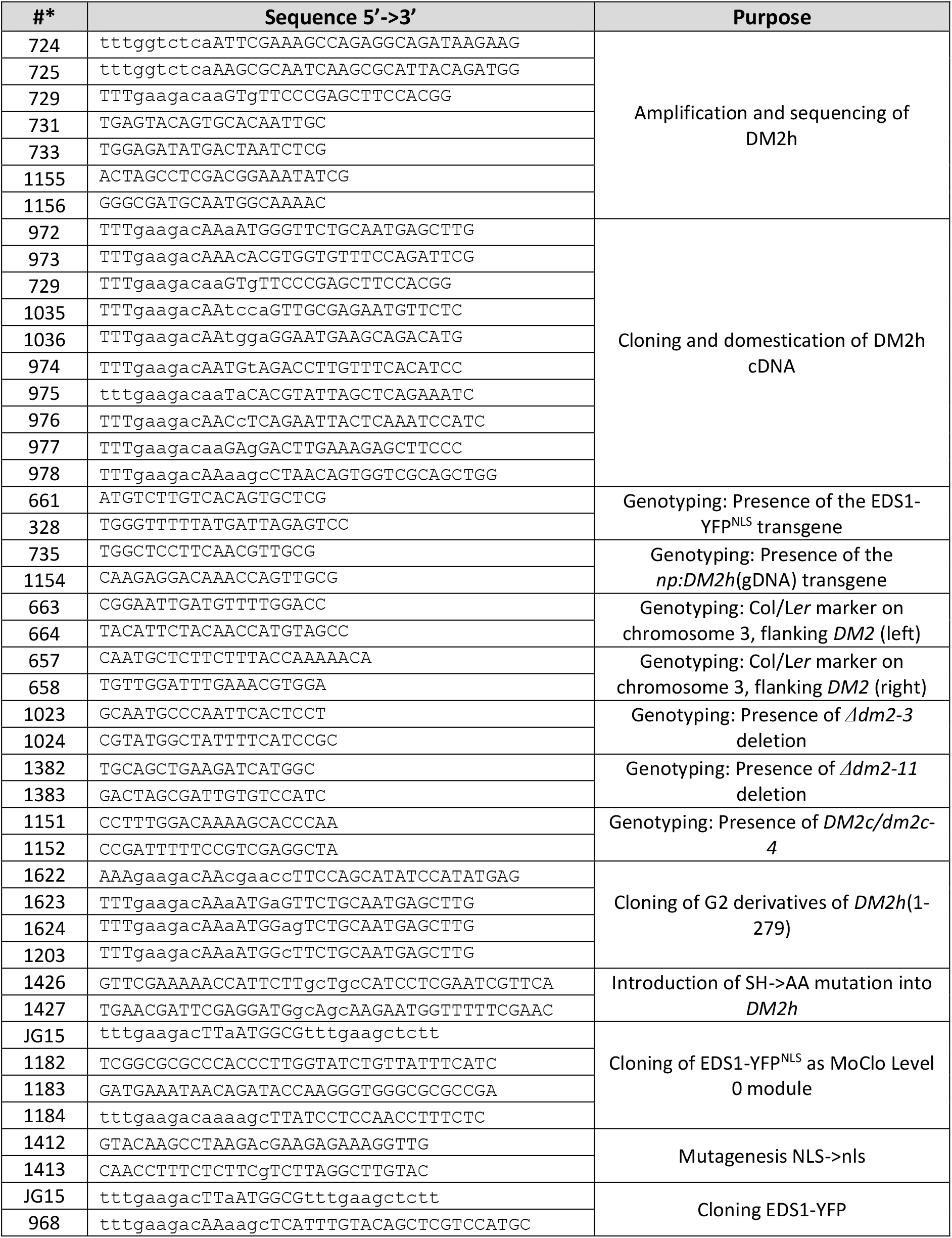
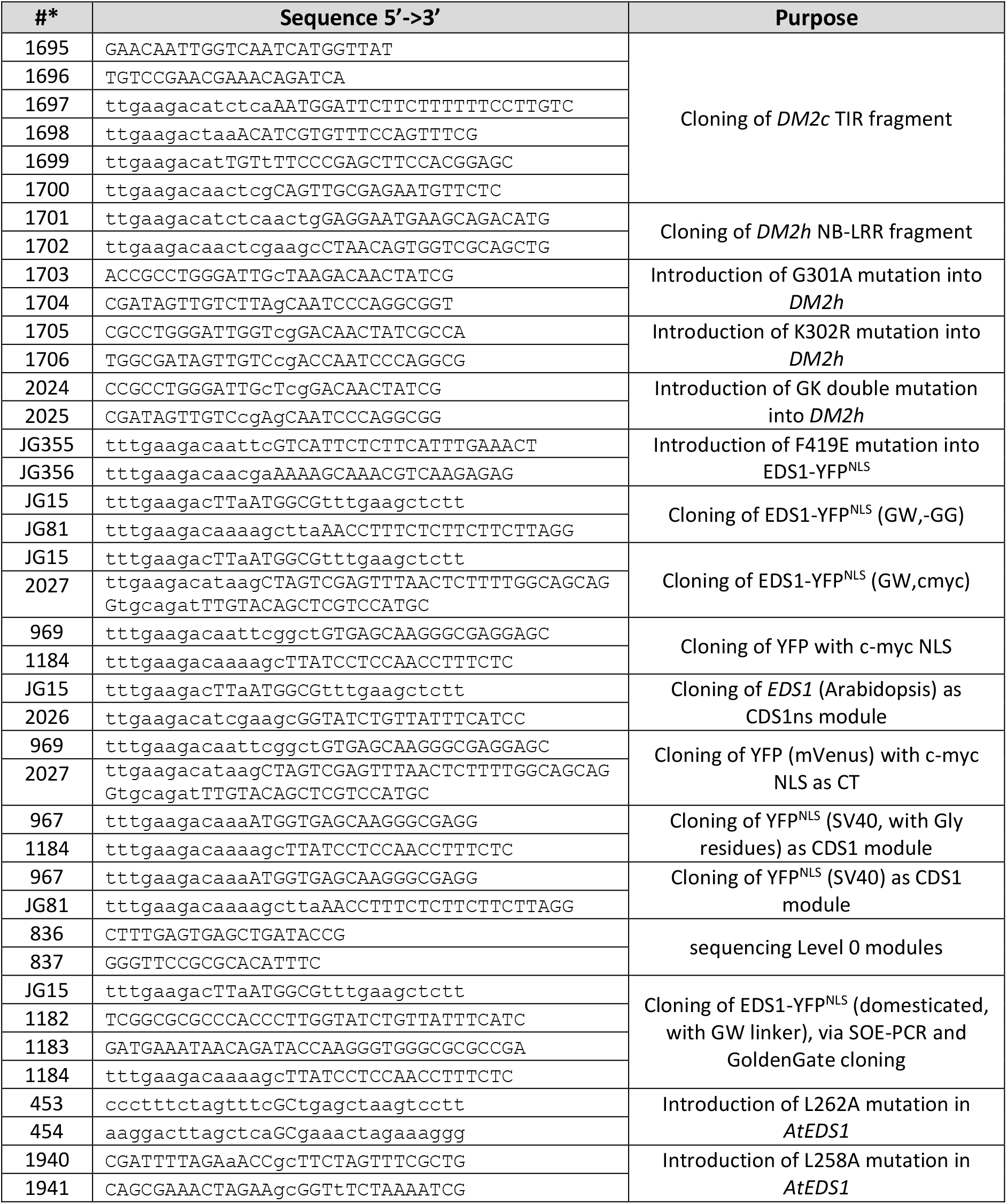
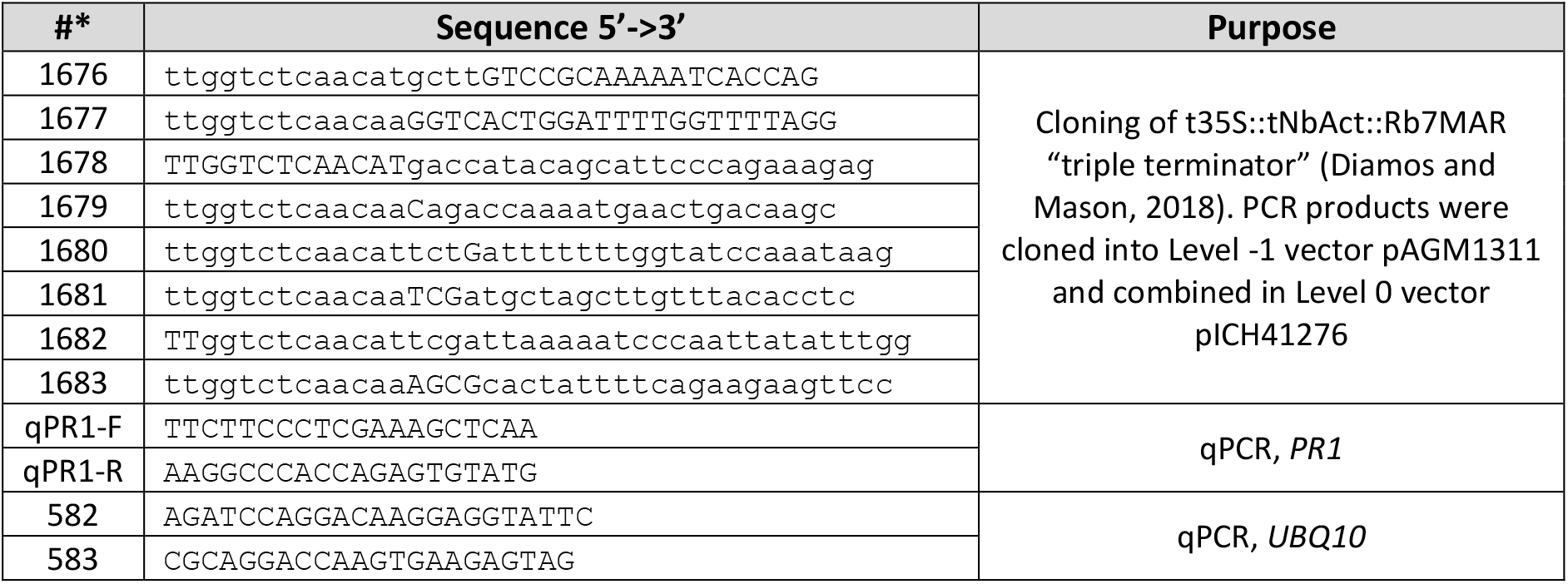
Oligonucleotides used in this study

**Table S3:** Plasmids used in this study

**Previously reported plasmids:**
| name | Description | Reference |
| --- | --- | --- |
| pUC57-Bsal | pUC57 derivative without *Bsal* site in *bla* | (Ordon et al., 2017) |
| pVM_BGW | T-DNA vector, pVS1 origin of replication |  |
| pJOG899 | DM2h<sub>(1-279)</sub> without STOP in pAGM1287 | (Gantner et al., 2018;<br>Gantner et al., 2019) |
| pJOG604 | FAST as 1-1r |  |
| pJOG640 | [Pro+5Uf] p35S-long |  |
| pJOG176 | mEGFP [CT] |  |
| pJOG25 | pEDS1 [Pro+5U] |  |
| pAGM9121 | Level 0 vectors Modular Cloning system | (Engler et al., 2014) |
| pICH41421 |  |  |
| pICH41308 |  |  |
| pAGM41276 |  |  |
| pICH41295 |  |  |
| pAGM1287 |  |  |
| pICH47732 (1-1f) | Level 1 vectors Modular Cloning system |  |
| pICH47742 (1-2f) |  |  |
| pICH47751 (1-3f) |  |  |
| pICH51266 | 35S Promoter [Pro+5U] |  |
| pICH51277 | 35S Promoter [Pro+5U] |  |
| pICH41414 | 35S terminator [3U+ter] |  |
| pICH50881 | end linker |  |
| pICH41744 | end linker |  |
| pICSL30008 | 6xHA [NT] |  |
| pAGM1311 | Level -1 universal acceptor |  |
| pXCSG-YFP | pAM-PAT-based GW destination vector;<br>35S:GW-YFP_t35S | pAM-PAT -> GenBank<br>AY436765.1 |

| Name | Recipient | Description | Oligo-nucleotides* |
| --- | --- | --- | --- |
| pJOG900 | pAGM1287 | DM2h<sub>(1-279)</sub> [G2S] without STOP | JS1623/1622 |
| pJOG901 | pAGM1287 | DM2h<sub>(1-279)</sub> [G2E] without STOP | JS1624/1622 |
| pJOG908 | pAGM1287 | DM2h<sub>(1-279)</sub> [G2A] without STOP | JS1203/1622 |
| pJOG941 | pAGM1287 | DM2h<sub>(1-279)</sub> [SH-AA] without STOP | SDM JS1426/27 |
| pJOG273 | pICH41308 | cDNA of DM2h as CDS1 (domesticated) | see methods |
| pJOG486 | pICH41308 | EDS1-YFP<sup>NLS</sup> (domesticated, with GW linker) | JG15/JS1182,<br>JS1183/1184 |
| pJOG813 | pICH41308 | EDS1-YFP<sup>NLS</sup> [F419E] | JG355/56, *DpnI*-<br>*BpiI*+ligase |
| pJOG1165 | pICH41308 | EDS1-YFP<sup>NLS</sup> [F419E,L262A] | SDM JS453/454 |
| pJOG1166 | pICH41308 | EDS1-YFP<sup>NLS</sup> [F419E,L258A,L262A] | JS1940/41 |
| pJOG1008 | pICH41276 | t35S::tNbAct::Rb7MAR terminator (Diamos and Mason, 2018) as MoClo Level 0 module<br>[intermediate cloning of PCR products into Level -1 vector pAGM1311] | JS1676/1677,<br>JS1678/1679,<br>JS1680/1681,<br>JS1682/1683 |
| pJOG747 | pICH41308 | EDS1-YFP (including GW linker) as CDS1 module | JG15, JS968 |
| pJOG993 | pAGM9121 | DM2c_TIR | JS1697/98, JS1699/1700 on an amplicon generated with JS1695/96 on gDNA |
| pJOG994 | pAGM9121 | DM2h_NB-LRR | JS1701/1702 |
| pJOG996 | pICH41308 | DM2h[G301A] as CDS1 | SDM 1703/04 |
| pJOG1011 | pICH41308 | DM2h[K302R] as CDS1 | SDM1705/06 |
| pJOG1213 | pICH41308 | DM2h[GK-AR] as CDS1 | SDM JS2024/25 |
| pJOG1312 | pICH41308 | EDS1-YFP<sup>NLS</sup> (GW,-GG) | JG15/JG81 |
| pJOG1337 | pICH41308 | EDS1-YFP<sup>NLS</sup> (GW,cmcy) | JG15/JS2027 |
| pJOG1329 | pAGM1301 | YFP<sup>NLS</sup> as CT (without GG) | JS969/1184 |
| pJOG1212 | pAGM1287 | EDS1 as CDS1ns module | JG15/JS2026 |
| pJOG1235 | pAGM1301 | YFP with c-myc NLS as CT | JS969/2027 |
| pJOG822 | pICH41308 | YFP<sup>NLS</sup>(SV40, with GG) as CDS1 | JS967/1184 |
| pJOG1311 | pICH41308 | YFP<sup>NLS</sup> (SV40) as CDS1 | JS967/JG81 |

\* sequences of oligonucleotides are provided in Table S2.

Level 1 modules (assembled as described in Engler et al., 2014):
| Name | Recipient | Inserts | Description |
| --- | --- | --- | --- |
| pJOG999 | pICH47742 | pICH51277, pICH41414, pJOG176, pJOG899 | 35S:DM2h<sub>(1-279)</sub>-GFP |
| pJOG1000 | pICH47742 | pICH51277, pICH41414, pJOG176, pJOG900 | 35S:DM2h<sub>(1-279)</sub>[G2S]-GFP |
| pJOG1001 | pICH47742 | pICH51277, pICH41414, pJOG176, pJOG901 | 35S:DM2h<sub>(1-279)</sub>[G2E]-GFP |
| pJOG1076 | pICH47742 | pICH51277, pICH41414, pJOG176, pJOG908 | 35S:DM2h<sub>(1-279)</sub>[G2A]-GFP |
| pJOG1077 | pICH47742 | pICH51277, pICH41414, pJOG176, pJOG941 | 35S:DM2h<sub>(1-279)</sub>[SH-AA]-GFP |
| pJOG274 | pICH47742 | pICH41414, pJOG273, pICH51277 | 35S:DM2h (cDNA) |
| pJOG546 | pICH47751 | pJOG486, pICH41414, pICH51277 | 35S:EDS1-YFP<sup>NLS</sup> |
| pJOG750 | pICH47751 | pJOG747, pICH41414, pICH51277 | 35S:EDS1-YFP |
| pJOG995 | pICH47742 | pICH51266, pICH41414, pJOG993, pJOG994 | 35S:DM2c(TIR)-DM2h(NB-LRR) |
| pJOG1009 | pICH47742 | pICH41414, pICH51277, pJOG996 | 35S:DM2h[G301A] |
| pJOG1194 | pICH47742 | pICH41414, pICH51277, pJOG1011 | 35S:DM2h[K302R] |
| pJOG1256 | pICH47742 | pICH41414, pICH51277, pJOG1213 | 35S:DM2h[GK->AR] |
| pJOG857 | pICH47742 | pJOG273, pICH41414, pICSL30008, pJOG640 | 35S:6xHA-DM2h |
| pJOG1010 | pICH47742 | pICH41414, pJOG996, pICSL30008, pJOG640 | 35S:6xHA-DM2h[G301A] |
| pJOG1195 | pICH47742 | pICH41414, pJOG1011, pICSL30008, pJOG640 | 35S:6xHA-DM2h[K302R] |
| pJOG1257 | pICH47742 | pICH41414, pJOG1213, pICSL30008, pJOG640 | 35S:6xHA-DM2h[GK-AR] |
| pJOG746 | pICH47751 | n/a; SDM on pJOG546, oligos 1412/ 1413 | 35S:EDS1-YFP<sup>NLS</sup> |
| pJOG814 | pICH47751 | pJOG813, pICH41414, pICH51277 | 35S:eds1(F419E)-YFP<sup>NLS</sup> |
| pJOG1167 | pICH47742 | pJOG25, pJOG486, pJOG1008 | pEDS1:EDS1-YFP<sup>NLS</sup> |
| pJOG1168 | pICH47742 | pJOG25, pJOG1166, pJOG1008 | pEDS1:EDS1-YFP<sup>NLS</sup>(F419,L258,L262) |
| pJOG1314 | pICH47751 | pICH51277, pICH41414, pJOG1312 | p35S:EDS1-YFP<sup>NLS</sup> (GW,-GG) |
| pJOG1338 | pICH47751 | pICH51277, pICH41414, pJOG1337 | p35S:EDS1-YFP<sup>NLS</sup>(GW,cmcy) |
| pJOG1330 | pICH47751 | pICH51277, pICH41414, pJOG1329,<br>pJOG1212 | p35S:EDS1-YFP<sup>NLS</sup> (SV40,SA) |
| pJOG1237 | pICH47742 | pICH51266, pJOG1212, pJOG1235,<br>pJOG1008 | p35S:EDS1-YFP<sup>NLS</sup>(cmcy,SA) |
| pJOG825 | pICH47751 | pJOG822, pICH41414, pICH51277 | p35S:YFP<sup>NLS</sup> (+GG) |
| pJOG1313 | pICH47751 | pICH51277, pICH41414, pJOG1311 | p35S:YFP<sup>NLS</sup> (-GG) |
| pJOG1350 | pICH47742 | pJOG1008, pJOG25, pJOG486 | pEDS1:AtEDS1-YFP<sup>NLS</sup> (GW, SV40) |
| pJOG1351 | pICH47742 | pJOG1008, pJOG25, pJOG1337 | pEDS1:AtEDS1-YFP<sup>NLS</sup> (GW, cmcy) |
| pJOG1352 | pICH47742 | pJOG1008, pJOG25, pJOG1212,<br>pJOG1329 | pEDS1:AtEDS1-YFP<sup>NLS</sup> (SA, SV40) |
| pJOG1353 | pICH47742 | pJOG1008, pJOG25, pJOG1212,<br>pJOG1235 | pEDS1:AtEDS1-YFP<sup>NLS</sup> (SA, cmcy) |

**Level 2 vectors (plant transformation vectors; assembled as described in Engler et al., 2014):**
| Name | Recipient | Inserts | Description |
| --- | --- | --- | --- |
| pJOG1169 | pJOG1051 | pJOG604, pJOG1167, pICH50881 | FAST-pEDS1:EDS1-YFP<sup>NLS</sup> |
| pJOG1170 | pJOG1051 | pJOG604, pJOG1168, pICH50881 | FAST-pEDS1:EDS1-YFP<sup>NLS</sup>(F419,L258,L262) |
| pJOG1354 | pJOG1051 | pJOG604, pICH41744, pJOG1350 | FAST-pEDS1:EDS1-YFP<sup>NLS</sup>(SV40) [GW] |
| pJOG1355 | pJOG1051 | pJOG604, pICH41744, pJOG1351 | FAST-pEDS1:EDS1-YFP<sup>NLS</sup>(cmcy) [GW] |
| pJOG1356 | pJOG1051 | pJOG604, pICH41744, pJOG1352 | FAST-pEDS1:EDS1-YFP<sup>NLS</sup>(SV40) [SA] |
| pJOG1357 | pJOG1051 | pJOG604, pICH41744, pJOG1353 | FAST-pEDS1:EDS1-YFP<sup>NLS</sup>(cmcy) [SA] |

**Other plasmids**
| Name | Description |
| --- | --- |
| pJOG1051 | Level 2 recipient prepared from pVM-Bpil; contains ccdB (excised during cloning) |
| pCW5 | 35S:DM2h (gDNA) |
| pAM-PAT<br>GW OLD3 | 35S:OLD3; OLD3 cDNA recombined into pAM-PAT GW by LR reaction |
| pAM-PAT<br>GW old3-1 | 35S:old3-1; old3-1 cDNA recombined into pAM-PAT GW by LR recombination. |
| pJOG802 | SRF3(Kondara) in pJOG130 (Gateway entry) |
| pJOG826 | SRF3 in pAM-PAT GW, created by LR reaction |
| pXCSG-YFP<br>DM2h | gDM2h (noSTOP) in pXCSG-YFP, created by LR reaction |

**Supplemental Figure 1:**
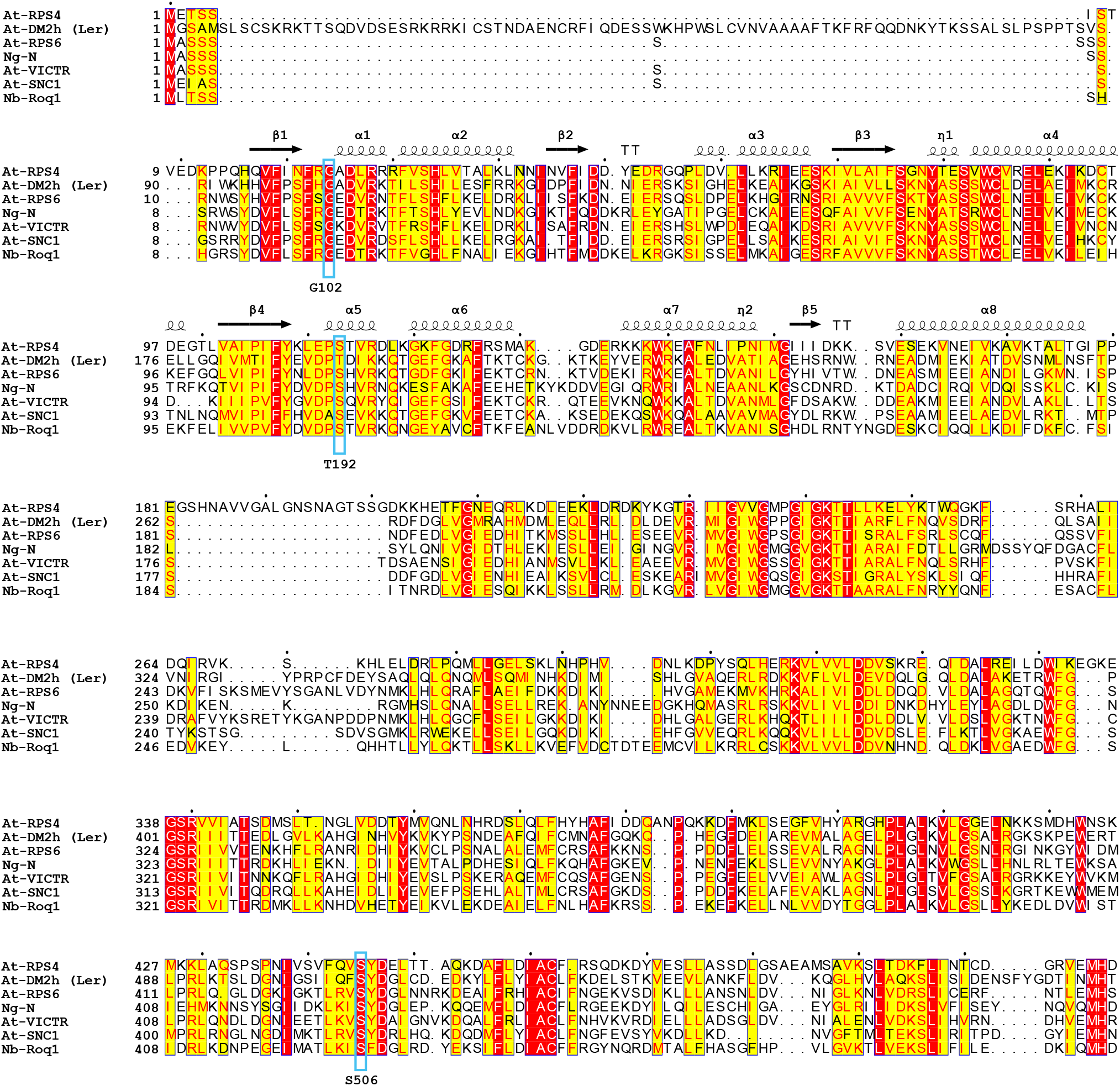
Sequence alignment of TIR and NB -ARC domains. Indicated proteins were aligned using tcoffee, and the alignment was rendered using ESPript. Secondary structure elements of the Arabidopsis RPS4 TIR domain (PDB:4C6R) are shown above the alignment. Positions / amino acids affected by *nde* mutations are boxed. At - *Arabidopsis thaliana*, Ng - *Nicotiana glutinosa*, Nb - *Nicotiana benthamiana*. UniProt identifiers: AtRPS4 - Q9XGM3, AtRPS6 - P0DKH6, NgN - Q40392, AtVICTR - F4KHI3, AtSNC1 - O23530, NbRoq1 - A0A290U7C4.

**Supplemental Figure 2:**
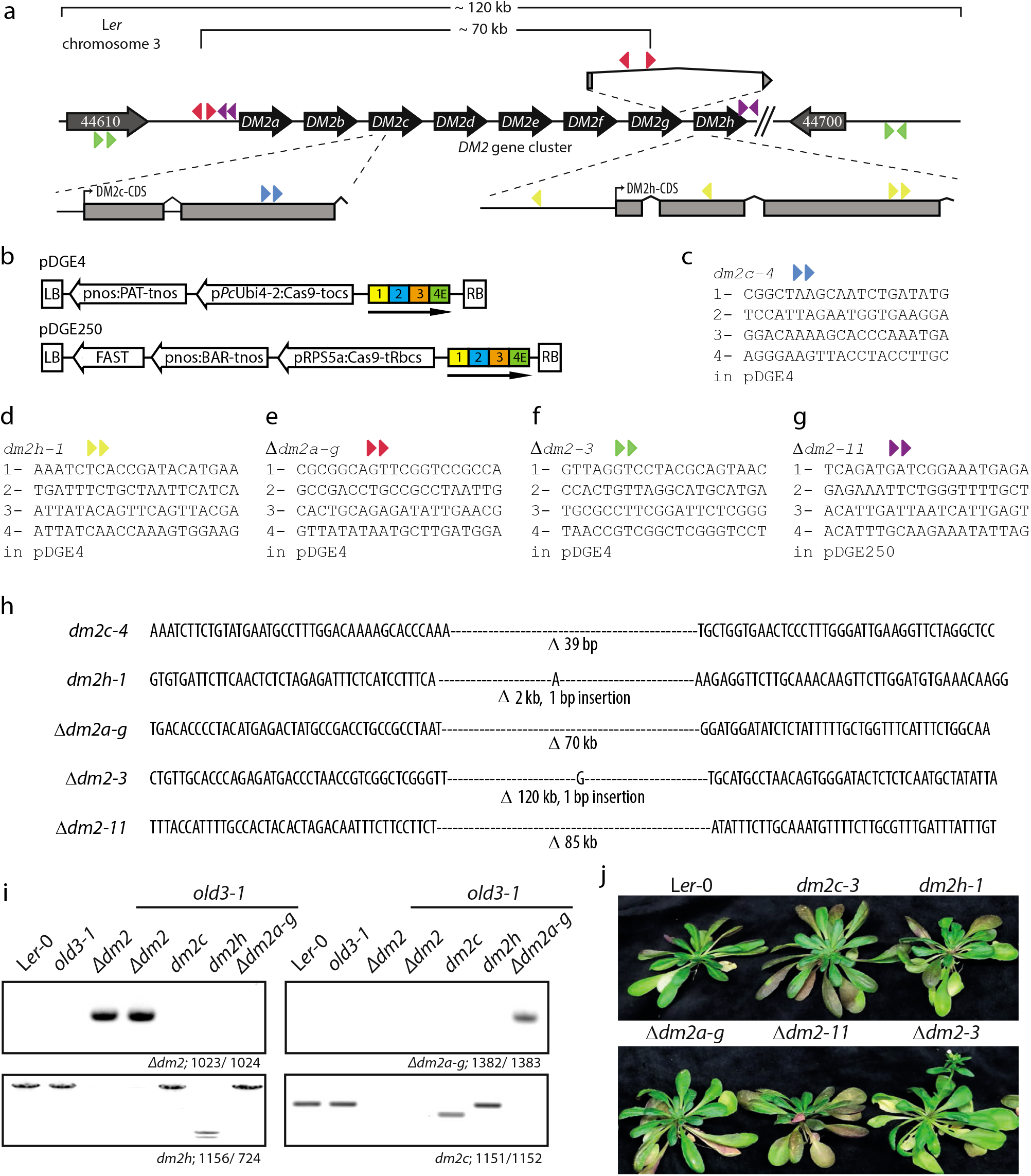
*dm2* mutant lines used in this study. a) Schematic drawing of the *DM2^Ler^* region with sgRNA target sites (colored triangles) indicated. b) Schematic drawing of constructs (T-DNA region) used for genome editing. c) - g) Sites targeted by RNA-guided nucleases for generation of indicated mutant lines. Color code corresponds to panel a). h) Precise lesions detected in *dm2* mutant lines used in this study. i) Genotyping of *dm2* mutant lines. j) Early flowering phenotype of *Δdm2-3* mutant plants. Plants were grown side-by-side for 7 - 8 weeks under short day conditions. Representative plants were selected for documentation.

**Supplemental Figure 3:**
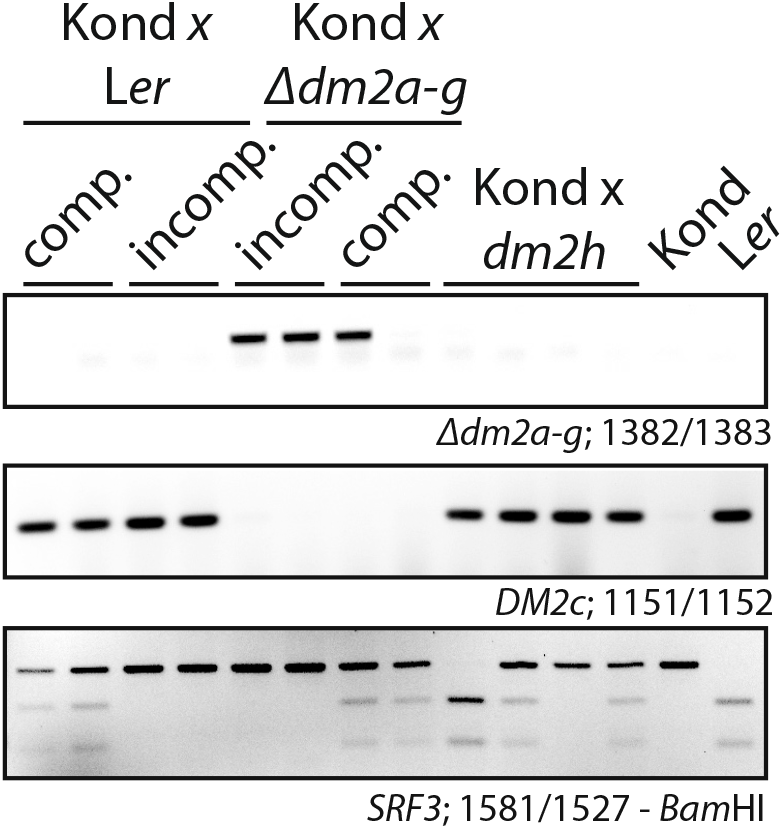
Genotyping of hybrids obtained from crosses to accession Kondara. Compatible (comp.) and incompatible (incomp.) hybrids obtained from F_2_ generations from crosses of L*er* or *dm2* mutant lines to Kondara, as shown in Figure 4, were genotyped by PCR. Primer combination 1382/1383 detects the *Δdm2a-g* deletion. Primer combination 1151/1152 detects presence of *DM2c*, and was used to probe presence/absence of the *DM2* cluster. A CAPS marker differentiating *SRF3* from L*er* and Kondara shows segregation of these alleles in compatible hybrids, while only the SRF3^Kond^ allele was detected in incompatible hybrids, as expected.

**Supplemental Figure 4:**
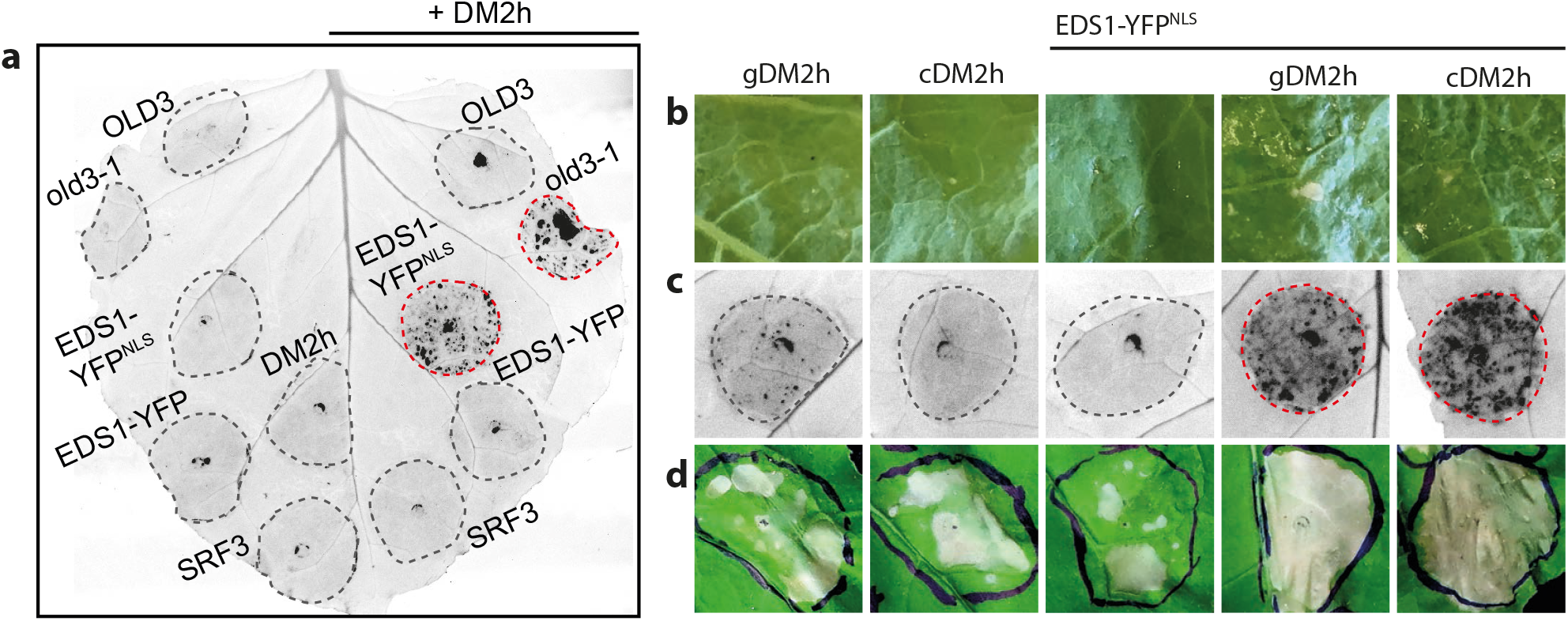
A transient assay for reconstitution of DM2h activation in *N. benthamiana* based on protein co-expression and cell death induction. a) Induction of HR-like cell death upon co-expression (by Agroinfiltration) of DM2h (35S:gDM2h) with different autoimmunity inducers or variants thereof. Agrobacterium strains for the expression of indicated proteins (under 35S promoter control) were infiltrated into *Nb* wild-type leaves. The formation of HR-like cell death was photographed 7dpi on a a gel documentation system. b) - d) as in a), but the DM2h gDNA and a cDNA construct (corresponding to the gene model shown in Figure 1a) were compared for cell death induction, and different methods were used to document plant reactions. b) shows macroscopically visible plant reactions. Tissues from co-expression became glossy at the surface, and small necrotic patches appeared. c) shows cell death reactions as visualized under UV light using a gel documentation system. d) shows macroscopic plant reactions at 5 dpi when plants were incubated for 2d in the dark subsequent to infiltration to enhance cell death induction, as described before (Chae et al., 2014; Qi et al., 2018). Similar results were obtained when expressing DM2h from a gDNA or a cDNA construct. cDNA constructs were used for site-directed mutagenesis and all further experiments.

**Supplemental Figure 5:**
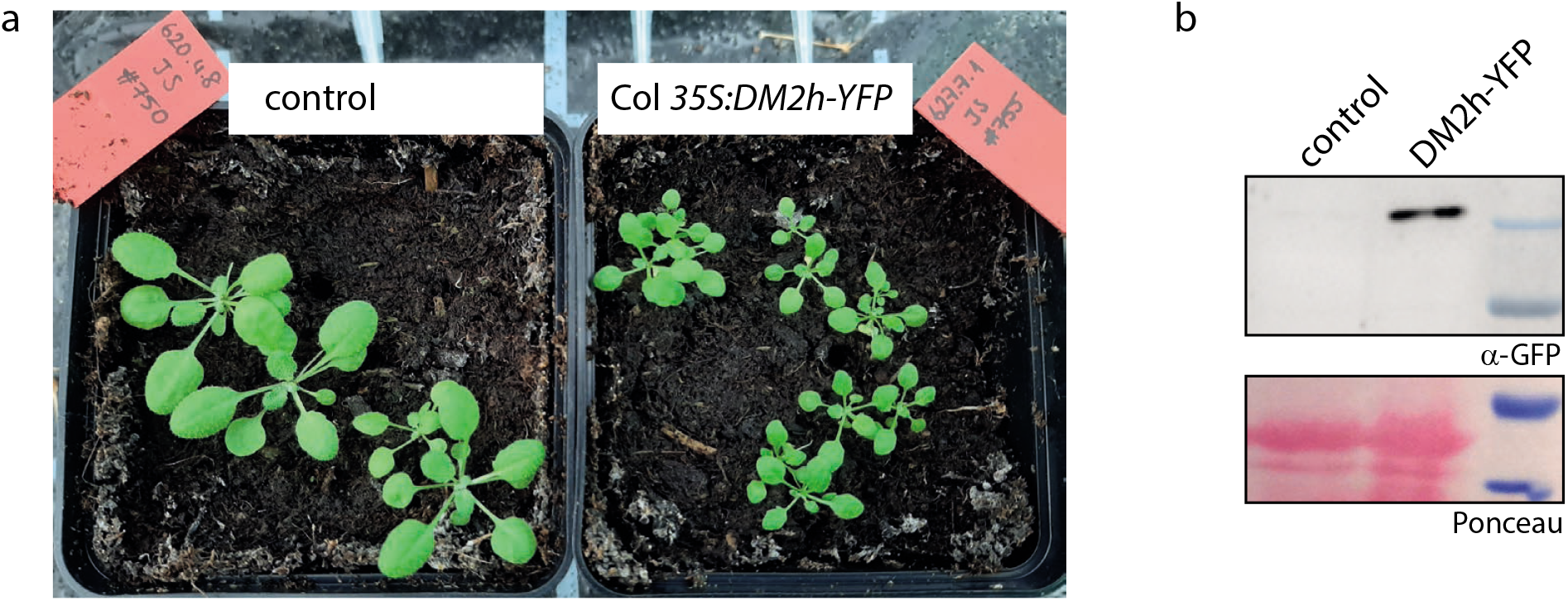
Growth phenotype and immunoblot analysis of DM2h-YFP transgenic plants. a) Homozygous T_3_ transgenics containing *35S:gDM2h-YFP* and a control line were grown side-by-side under short day conditions for four weeks. DM2h-YFP transgenic plants were smaller, and showed some necrotic lesions indicative of autoimmunity on lower leaves, suggesting that the DM2h-YFP fusion protein is at least partially functional. b) Immunodetection of the DM2h-YFP fusion protein. Tissue samples were taken from the same plants that were used for live cell imaging (Figure 5g).

**Supplemental Figure 6:**
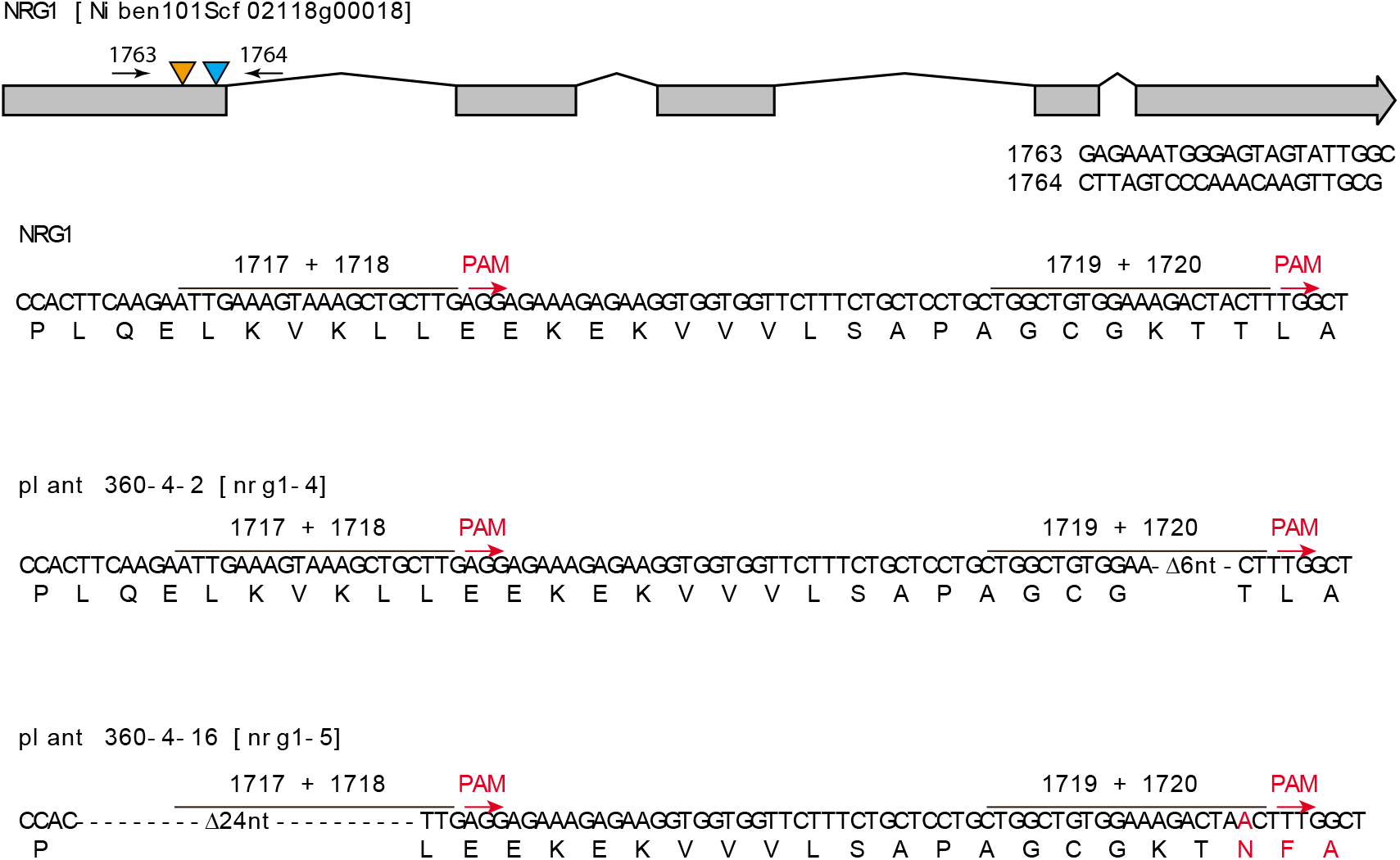
Molecular details on *Nbnrg1* mutant lines generated in this study. The *NbNRG1* locus was targeted for genome editing using sgRNAs previously reported in Qi et al. (2018). The locus with the relative position of the target sites is show as an overview and at the nucleotide level. Two different homozygous lines, *nrgl-4* and *nrg1-5*, were isolated from a segregating T_1_, population (plant 360-4). PCR primers used for screening and sequencing of alleles (1763/1764) are indicated. Final lines do not contain the T-DNA anymore. Similar phenotypes were observed with both lines.

**Supplemental Figure 7:**
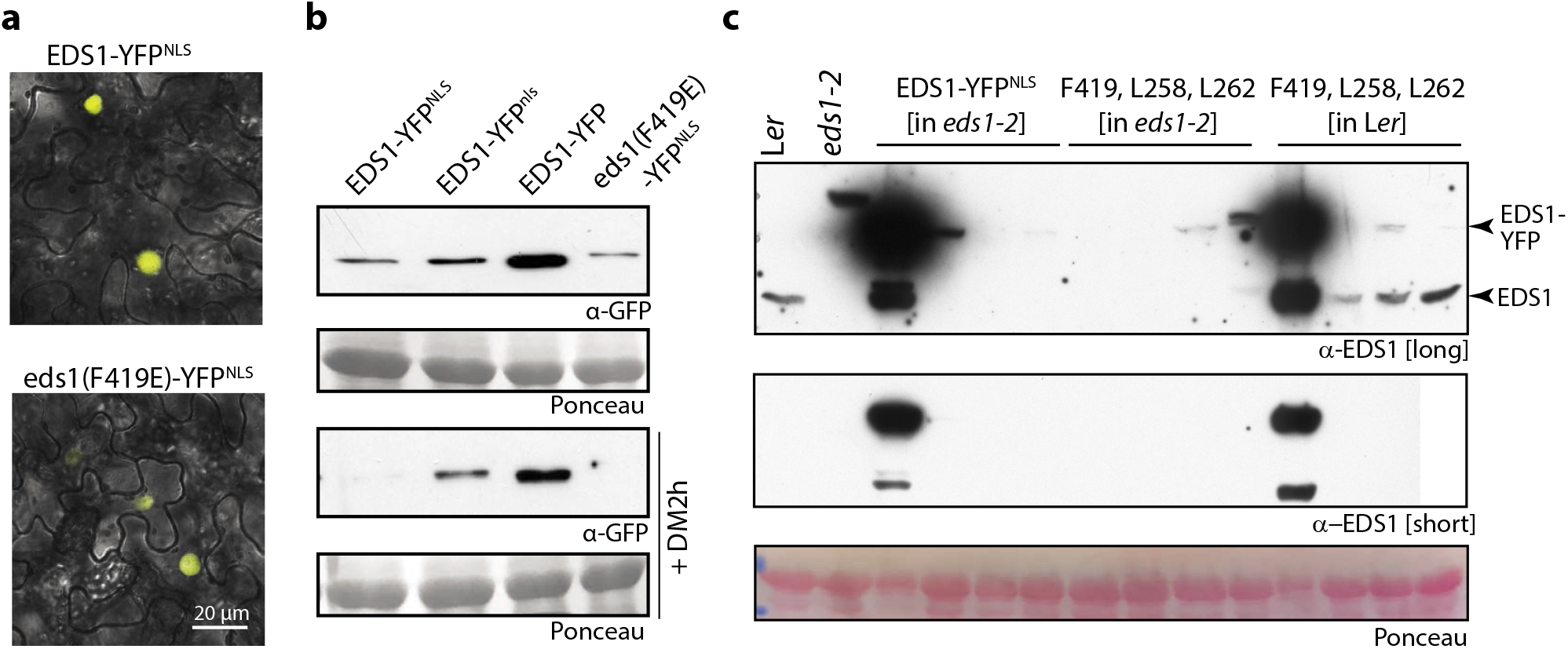
Immunodetection and subcellular localization of EDS1-YFP^NLS^ variants differing in immune signaling competency. a) Localization of EDS1-YFP^NLS^ variants In *N. benthamiana* (in co-expression with DM2h) as assessed by confocal laser scanning microscopy at 3 dpi. An overlay of bright field and YFP imaging (yellow) is shown. Similar localizations were observed when EDS1 variants were expressed without DM2h. Scale bar = 20 μm. b) Immunodetection of fusion proteins expressed in Figure 6a and S7a. Tissues were harvested 3 dpi, and total protein extracts used for SDS-PAGE and immunodetection. c) Immunodetection of EDS1-YFP^NLS^ variants in primary Arabidopsis transformants (in support of Figure 6b). Transgenic seeds were selected by FAST seed fluorescence. Plants were grown 7d under short day conditions, 20d under immune suppressive high temperature conditions and shifted to 16/14°C for 8d. Lanes with strong EDS1-YFP signals correspond to autoimmune plants. Remaining plants were selected randomly. Low accumulation of the EDS1-YFP^NLS^ [F419E, L258A, L262A] in the *eds1-2* mutant background is most likely due to low *EDS1* expression in absence of

**Supplemental Figure 8:**
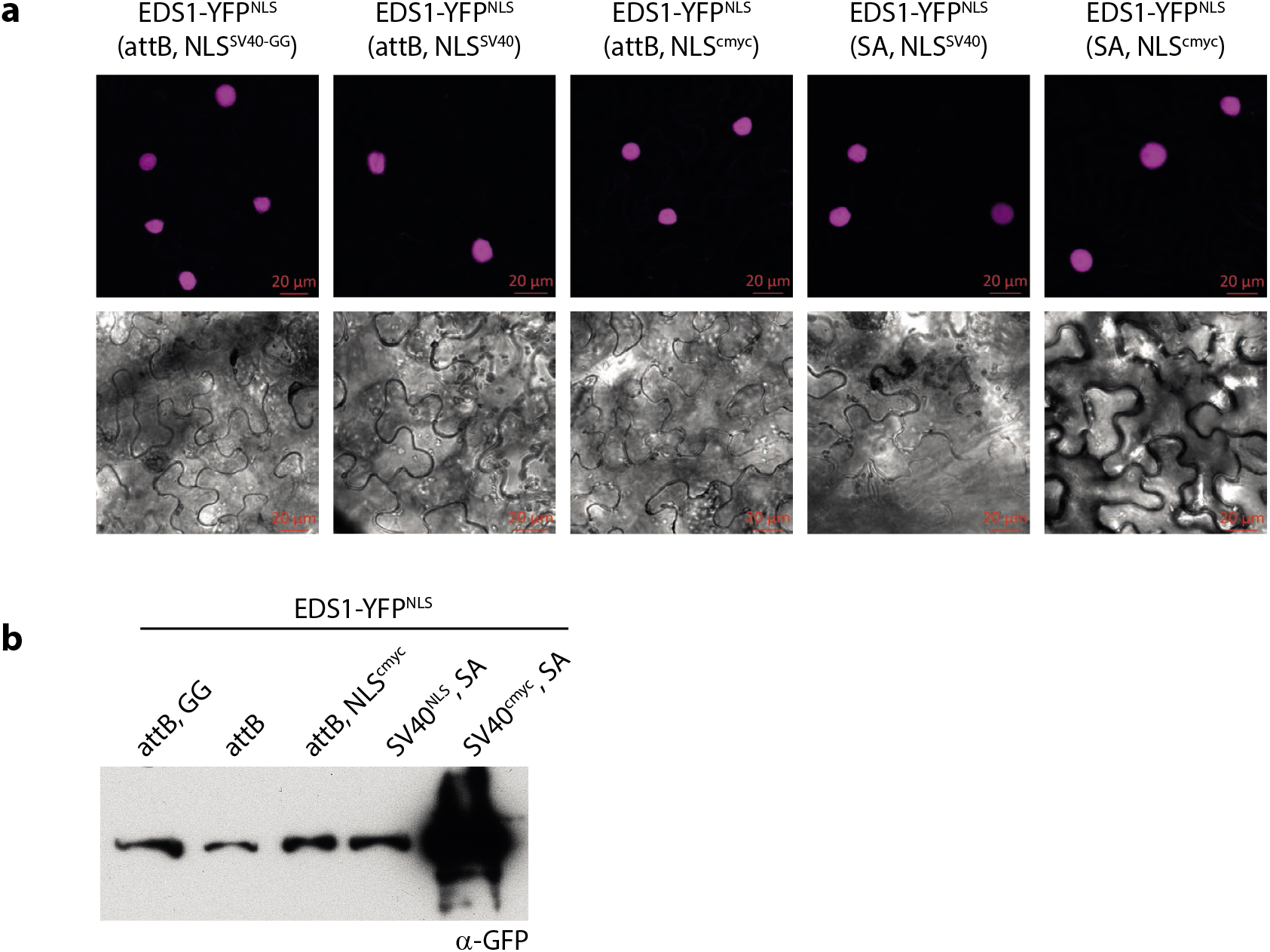
Immunodetection and subcellular localization of EDS1-YFP^NLS^ variants differing in NLS and linker sequences. a) Localization of EDS1-YFP^NLS^ variants (in co-expression with DM2h) as assessed by confocal laser scanning microscopy at 3 dpi. YFP channel (magenta) and bright field images are shown. Similar results were obtained upon co-expression of fusions without DM2h. Scale bar = 20 μm. b) Immunodetection of fusion proteins expressed in Figures 7 and S8a. Tissues were harvested 3 dpi from the same

## Acknowledgements

This work was funded by GRC grant STU 642-1/1 (Deutsche Forschungsgemeinschaft, DFG) and seed funding by the CRC 648 (DFG) to JS. UB is grateful for financial support by the Leibniz price from the DFG and the Alfried Krupp von Bohlen und Halbach Stiftung. FF was financed by an ERASMUS mobility program. We are grateful to Bianca Rosinsky for taking care of plant growth facilities and growing plants, and to Ruben Alcazar and Jane Parker for providing plasmids containing *SRF3*^Kond^ or *RPP1-like R8/DM2h* genomic DNA fragments. Jan Kemna is acknowledged for cloning of the DM2h cDNA, and Samuel Grimm for primary screening of DM2h-YFP Arabidopsis transformants.

